# Lifestyle-mediated shifts in the gut microbiome and host genetics modulate gene expression in intestinal organoids

**DOI:** 10.64898/2026.11.07.765058

**Authors:** Shreya Nirmalan, Sabrina Arif, Gabrielle Garlicki, Adnan Alazizi, Kaitlyn Hutchins, Mathilde Poyet, Mathieu Groussin, Peter Carbonetto, Roger Pique-Regi, Ran Blekhman, Francesca Luca

**Affiliations:** Center for Molecular Medicine and Genetics, Wayne State University, Detroit, MI; Section of Genetic Medicine, Department of Medicine, University of Chicago, Chicago, IL; Department of Human Genetics, University of Chicago, Chicago, IL, USA; Institute of Experimental Medicine, Kiel University, Kiel, Germany; Global Microbiome Conservancy, microbiomeconservancy.org; Department of Biological Engineering, Massachusetts Institute of Technology, Cambridge, MA; Institute of Clinical Molecular Biology, Kiel University, Kiel, Germany; Department of Obstetrics and Gynecology, Wayne State University, Detroit, MI

## Abstract

Industrialization-associated shifts in the gut microbiome have been linked to disease, but the mechanisms by which microbial communities influence host gene regulation remain poorly understood. We developed a live microbiome-colonic organoid (colonoid) co-culture system to investigate genotype-dependent host responses to gut microbial community variation. Colonoid 2D monolayers from healthy donors were co-cultured with live gut microbial communities from individuals living urban or rural lifestyles, followed by RNA-sequencing. Urban and rural microbiomes elicited largely shared transcriptional responses, however they induced distinct host genetic effects on gene expression. We identified 511 host genes with microbiome-dependent allele-specific expression (ASE) which were involved in epithelial architecture, cell-matrix interactions and structural integrity. Notably, 149 host genes had distinct allelic effects between Urban and Rural microbiomes (urban-rural condition specific ASE or cASE), revealing genotype-dependent host responses to lifestyle-associated microbiomes. Urban-rural cASE was associated with microbial community composition, predominantly involving *Prevotella* and *Bacteroides* species. Targeted *Prevotella copri* spike-in recapitulated *Prevotella*-associated allelic effects at specific loci, including one in the IBD-associated autophagy gene, *ATG4B*. Together, these findings establish an experimental framework to investigate genotype-by-microbiome interactions, and demonstrate that lifestyle-mediated shifts in the microbiome modify host *cis*-regulatory genetic effects on gene expression.

## Introduction

The gut microbiome, consisting of trillions of microorganisms that inhabit the gastrointestinal tract, forms a dynamic interface between the host and the external environment. Through modulating immune function, nutrient processing and metabolite production, the gut microbiome plays a critical role in maintaining host intestinal homeostasis^1,2^. Variation in gut microbiome composition is shaped by lifestyle factors including diet, medication use and geography^3–5^. One major source of such variation is urbanization, the emergence of densely populated settlements, which is associated with distinct shifts in microbiome composition such as reduced microbial diversity, enrichment of *Bacteroides* species and depletion of *Prevotella* compared to rural-associated microbiomes^6–11^. Many of these same lifestyle changes have accompanied the global rise in non-communicable diseases, including inflammatory bowel disease, obesity and type 2 diabetes^12–14^. Alterations in gut microbiome composition have also been implicated in the pathogenesis of these disorders^15^, motivating the need to understand the mechanisms through which shifts in microbiome composition associated with urbanization influences host biology and disease risk.

One mechanism through which microbiome composition may influence host health is modulating host gene regulation in the colon, where host cells are directly exposed to microbial signals. Previous studies have demonstrated that the gut microbiome is associated with differential gene expression in the intestinal epithelium under both healthy and disease states^16–18^. Our group previously established an *in vitro* experimental system to study gene expression response to live microbial communities in human colonic epithelial cells^19^. Manipulation of the microbiota composition was found to cause predictable changes in host gene expression, and that genes whose expression varies with microbiome composition are enriched for genes associated with IBD risk through genome wide association studies^20,21^. Using this co-culture system, we found that urban-associated microbiomes and rural-microbiomes induce distinct transcriptional responses in the host, with urban microbiomes modulating host innate immune responses^22^. Collectively, these studies demonstrate that variation in gut microbiome composition can alter host gene expression in the colonic epithelium.

Host genetic variation modulates host gene regulation, and *cis*-regulatory variants associated with inter-individual variation in gene expression are also associated with human disease and complex traits^23,24^. These *cis-*regulatory effects vary across environmental contexts, also known as Genotype x Environment interactions (GxE)^25^. Consistent with this conceptual framework, we previously identified several host loci that exhibited interactions between host genotype and microbiome composition on host gene expression in intestinal tissue from patients with inflammatory bowel disease and healthy controls^26^. Additionally, prior work by our group used our established co-culture system in one primary colonocyte cell line to identify instances of *cis-*regulatory effects that were dependent on microbiome exposure^19^. However, extending this approach to study host genetic variation across multiple donors and diverse microbiome communities remains experimentally challenging using conventional colonic epithelial cell lines. While these cell lines provide a physiologically relevant model of the colonic epithelium, there are only a few lines available from genetically diverse donors. This motivates the need for an experimental model that enables direct interrogation of the interactions between lifestyle-mediated variation in gut microbiome composition and host genetic variation.

Human colonic organoids (colonoids) provide a physiologically relevant model to investigate host-microbiome interactions in the context of host genetic variation. Derived from primary intestinal stem cells, colonoids maintain normal colonic physiology, cellular differentiation and, most importantly, host genetic stability^27^. Furthermore, differentiation of colonoids into epithelial monolayers can be used to study how colonic epithelial cells respond to the microbiome under controlled experimental conditions^28^. However, intestinal organoid experiments querying multiple genetically distinct individuals and microbiome compositions are labor-intensive and difficult to scale, making conventional population-scale eQTL mapping studies challenging. Allele-specific expression (ASE) provides a powerful alternative for interrogating *cis*-regulatory effects in smaller cohorts across a large number of environmental contexts^29^. Previous studies have successfully utilized ASE to characterize GxE across diverse cellular and environmental contexts, revealing regulatory loci not previously identified through eQTL mapping studies, including variants implicated in disease risk through genome-wide association studies (GWAS)^29–31^. The ability to detect GxE, even in individual samples, makes an ASE framework ideal for testing whether lifestyle-associated microbial communities interact with host genotype to directly modulate host gene regulation.

Building on our established live microbiome co-culture system^19–22^, here we have developed a primary human colonoid co-culture model to investigate how lifestyle-associated variation in gut microbiome composition interacts with host genetic variation to influence gene regulation. Differentiated colonoid monolayers from healthy donors were co-cultured with live microbial communities isolated from individuals living urban and rural lifestyles followed by bulk RNA-sequencing. We first characterized host transcriptional responses to microbiome exposure, and then we used ASE to characterize cis-regulatory effects and context-dependent cis-regulatory effects across diverse cellular and environmental contexts. This study design provides a framework for identifying and characterizing the effects of lifestyle-associated microbiome composition on host transcriptional and allelic responses in the host intestinal epithelium.

## Results

### Urban and Rural Microbiomes Induce Shared Host Transcriptional Responses

To study the host responses to lifestyle-associated differences in gut microbiome composition, we used eight live microbial communities isolated from fecal samples from individuals living urban and rural lifestyles as part of the efforts of the Global Microbiome Conservancy (GMbC) including two urban-associated and two rural-associated microbiomes (Urban microbiomes and Rural microbiomes, respectively) from both Ghana and Rwanda, in order to minimize broader geographic and cultural confounding (**Fig 1A**). Donors of the rural microbiome samples lived subsistence-based farming lifestyles, while donors of the urban microbiome samples had previously transitioned from farming-based subsistence to more industrialized and urban lifestyles. This study design therefore allows the interrogation of host transcriptional shifts in response to microbiome variation arising from transitioning from a rural to urban lifestyle within a geographic region. To confirm that selected microbial communities were representative of microbiome compositional shifts associated with urban and rural lifestyles, we examined taxonomic profiles previously generated using shotgun metagenomic sequencing. Principal coordinates analysis revealed that PC1 explains 67.8% of the variation, and samples cluster along this PC based on lifestyle (urban/rural) (**Fig 1A**). At the genus level, rural microbiomes had higher abundance of *Prevotella* and lower abundance of *Bacteroides* and *Firmicutes* (one-sided Wilcoxon rank-sum test, *P* = 0.014 for each test), indicating that the selected samples captured characteristic lifestyle-associated variation observed in the larger GMbC cohort as well as other previous studies (**Fig 1C-E**)^6–11,22,32^.

**Figure 1.**
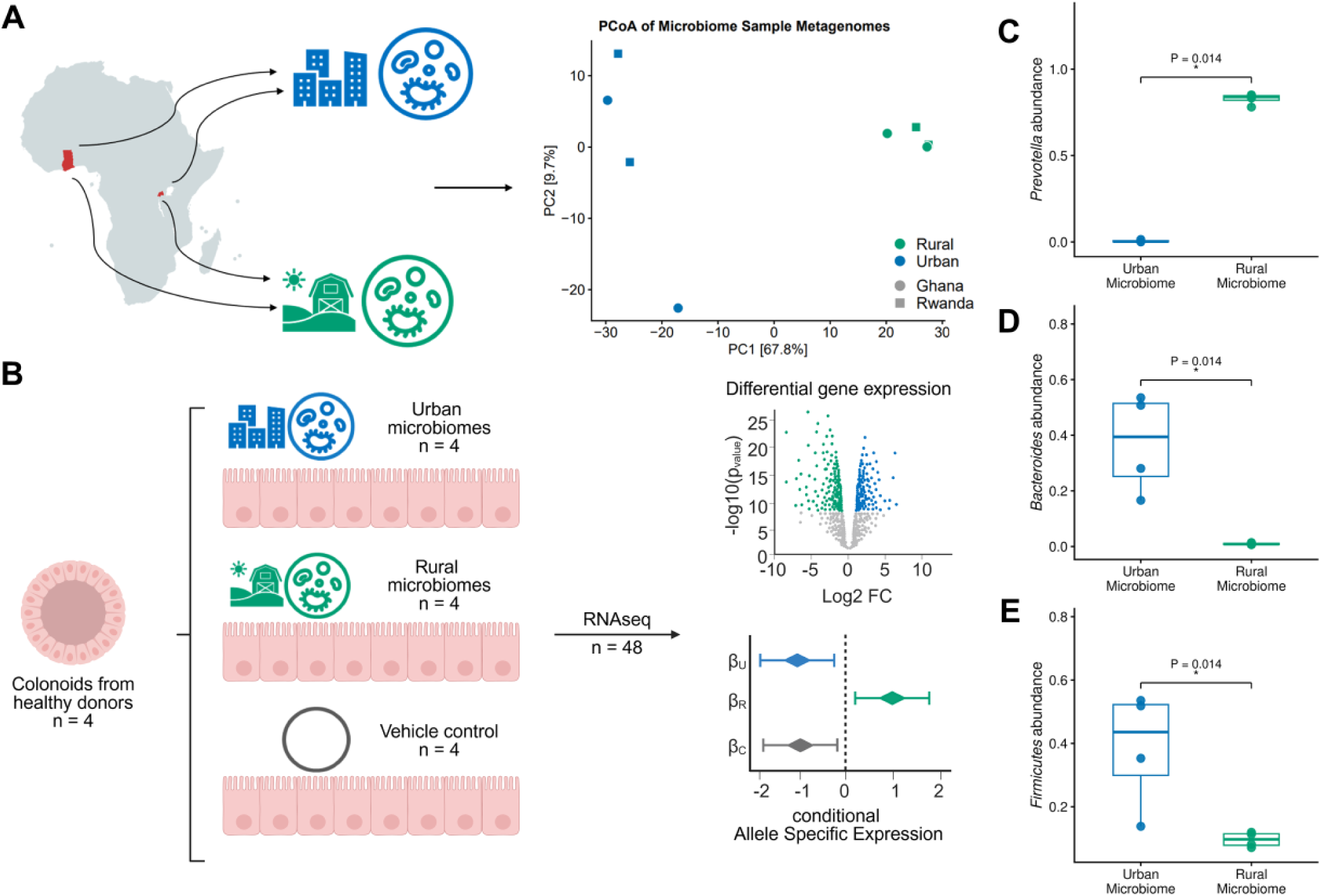
Study Design and Microbiome Sample Composition. **A)** Left: Live microbial communities originated from individuals from Ghana and Rwanda, with two Urban and two Rural microbiomes from each country. Right: Species-level principal coordinate analysis (PCoA) of microbiome samples. The x- and y-axes are the first and second principal components representing 67.8% and 9.7% of variability in microbiome composition, respectively. **B)** Colonoid monolayers from four healthy donors were co-cultured with four Urban or four Rural microbiomes or vehicle controls (n=4) and host transcriptional and allelic responses were measured using RNA sequencing (48 samples total).**C – E)** Boxplots of relative abundance of the genera *Prevotella, Bacteroides* and the phylum *Firmicutes* for the Urban (blue) and Rural (green) microbiomes used in the colonoid co-culture experiments. Points represent individual microbiomes and boxplots show the distribution across microbiomes. Differences between urban and rural microbiomes were assessed using one-sided Wilcoxon rank-sum tests, with exact *P* values shown above each comparison.

To investigate whether Urban and Rural microbiomes elicit distinct transcriptional responses, we differentiated colonoid monolayers from four healthy human donors and co-cultured them with each of the eight microbiomes or microbial-free media as the control, followed by RNA sequencing (**Fig 1B**). We observed considerable variation in the number of differentially expressed genes across colonoid lines in response to microbiome exposure, likely reflecting a substantial contribution of host genetics to the transcriptional response (Supplemental figure 1). When combining data across lines using a meta-analysis approach, we identified 823 host genes differentially expressed in response to microbiome exposure across all four lines (FDR < 10%) (**Fig 2A**), irrespective of the microbiome donor’s lifestyle. Gene ontology (GO) over representation analyses showed no significant enrichment for upregulated genes, while genes downregulated in response to microbiome exposure were primarily enriched for host responses to viral processes and type 1 interferon signaling (**Fig 2B**). The highest fold enrichment was observed for “cellular response to copper ion”, driven by the downregulation of five metallothionein isoforms (*MT1G, MT1A*, *MT1E, MT1H* and *MT1M*), a family of metal-binding proteins. In order to identify host transcriptional changes specific to Urban and Rural microbiomes, we characterized differential gene expression for the Urban microbiomes compared to control and for the Rural microbiomes compared to control, respectively. We identified a larger number of differentially expressed genes in response to Urban microbiomes compared to the Rural microbiomes (820 versus 387, respectively, FDR < 10%, **Fig 2C**), but changes in gene expression were significantly correlated between the two transcriptional responses (R = 0.56, p < 2 × 10^−16^). Consistent with this result, direct comparison of host transcriptional response to Urban versus Rural microbiomes identified only three differentially expressed genes (FDR < 10%): *DUOX2*, *GAPDH*, and *PPARG* (**Fig 2D-F**). Furthermore, GO overrepresentation analyses revealed that both Urban and Rural microbiomes led to downregulation of host viral response processes (**Supplemental Figure 2**) and upregulated host miRNA related biological processes (**Supplemental Tables 26, 28**). These results were further confirmed using multivariate adaptive shrinkage (MASH)^33^, which jointly analyzes the differential gene expression in the urban and rural conditions, relative to the control. From this combined analysis, we learned that the vast majority of genes show similar transcriptional responses to the Urban and Rural microbiomes compared to control, and large differences in these two transcriptional responses were rare; for example, only 51 genes showed differences in the responses to Rural and Urban microbiomes larger than a factor of 2, when considering the log-fold changes relative to control (**Supplemental Figure 3**). A much clearer trend was the tendency of the transcriptional response to be slightly but consistently larger in response to Urban microbiomes; specifically, among the 721 genes with significant response to both Urban and Rural microbiomes (lfsr < 0.1), on average the Urban log-fold changes were 1.17 fold larger than the Rural log-fold changes. Thus, despite the compositional differences in Urban and Rural microbiomes, they elicited largely similar host transcriptional responses with stronger effects induced by the Urban microbiomes.

**Figure 2.**
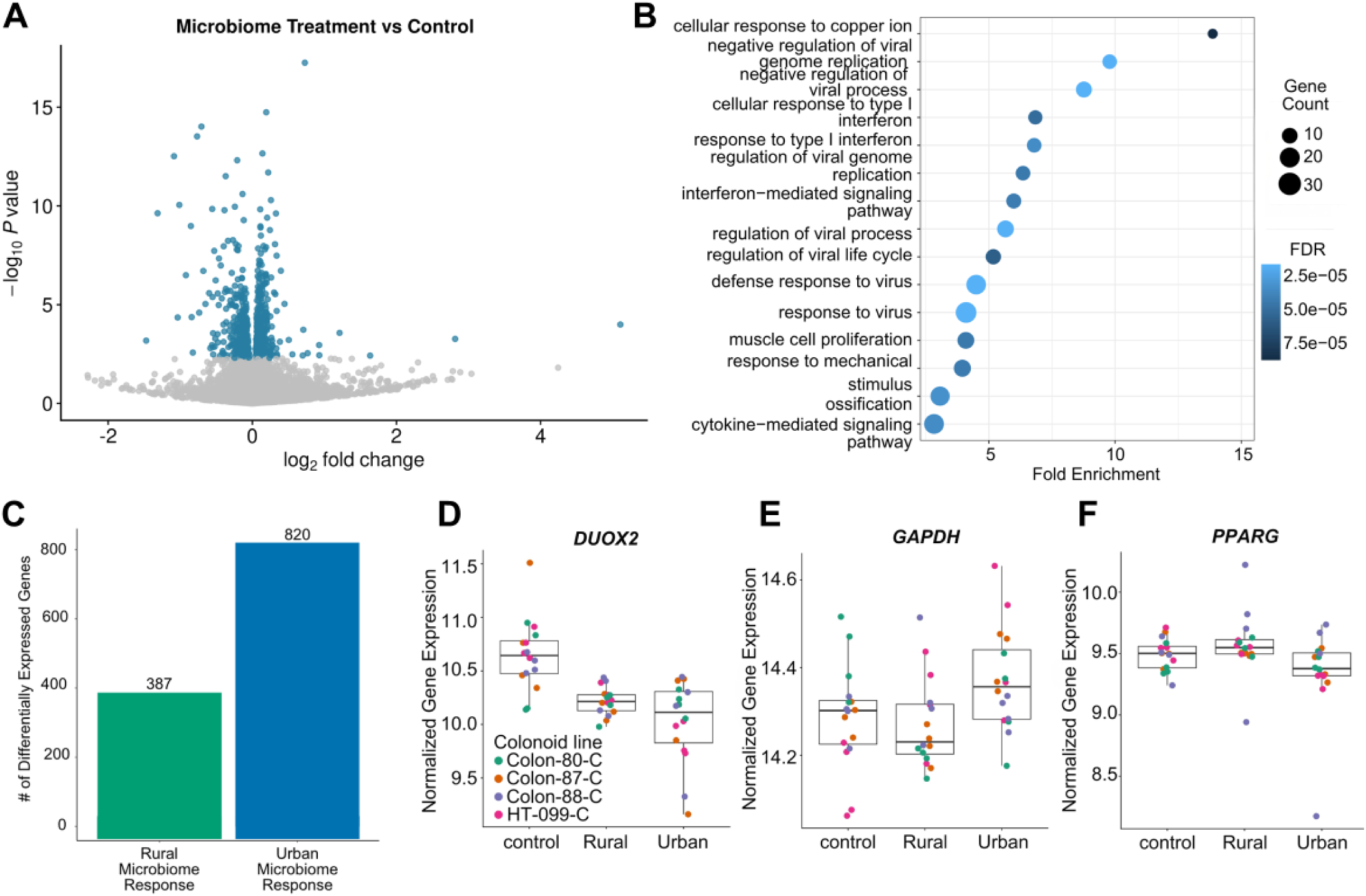
Urban and Rural microbiomes induce shared host transcriptional response. **A)** Volcano plot showing meta-analysis effect sizes for microbiome treatment versus controls. The x-axis represents the meta-analysis log2 fold change and the y-axis represents the –log10 p-value. Genes meeting the significance threshold (FDR < 10%) are shown in blue and non-significant genes are shown in gray. **B)** Dotplot of top 15 enriched Gene Ontology biological processes terms for genes downregulated in response to microbiome exposure (FDR < 10%). The Y-axis is the GO term and X-axis is the fold enrichment of genes associated with the GO term in differentially expressed genes. The size of the dot corresponds to the number of significant differentially expressed genes associated with the GO term and the color of the dot indicates the BH p-adjust value for the enrichment test. **C)** Barplot showing number of significantly differentially expressed genes in response to Urban or Rural microbiomes relative to control (FDR < 10%). **D-F)** Examples of genes differentially expressed between Urban and Rural microbiomes (FDR < 10%). Boxplots for each condition (Urban, Rural and control) are plotted separately with donor-line-corrected gene expression values plotted on the y-axis. Points are colored by which donor line the replicate/measurement originates from.

### Microbiome Exposure Reveals Widespread Shared and Context-Dependent Allele-Specific Expression

To investigate whether the microbiome modifies host genetics effects on gene expression, we first identified genes that exhibited allele-specific expression (ASE). ASE analysis compares the expression level of the two alleles at heterozygous loci in the coding sequence within the same individual (**Fig 3A**). In the absence of a genetic effect on gene expression, we expect similar levels of expression from each allele. An imbalance in allelic expression indicates the presence of a genetic effect on the expression of the gene. ASE allows for the characterization of genes with *cis*-regulatory variants while controlling for *trans*-effects and cellular environment. Differences in ASE across environmental conditions, i.e. condition-specific ASE (cASE), can identify context-dependent *cis*-regulatory effects reflective of GxE^29^. This framework allows for the identification of host genes that harbor interactions between host genetic variation and microbiome conditions, i.e., genotype x microbiome (GxM). We used MASH^33^ to jointly model ASE measured in different conditions and infer shared and condition-specific ASE. Our MASH analysis identified 8,252 SNPs in 3,510 genes with shared ASE, 595 SNPs in 511 genes with microbiome cASE, and 155 SNPs in 149 genes with Urban/Rural cASE (**Fig 3A**, **Supplemental Figure 4, Supplemental Table 31**).

**Figure 3.**
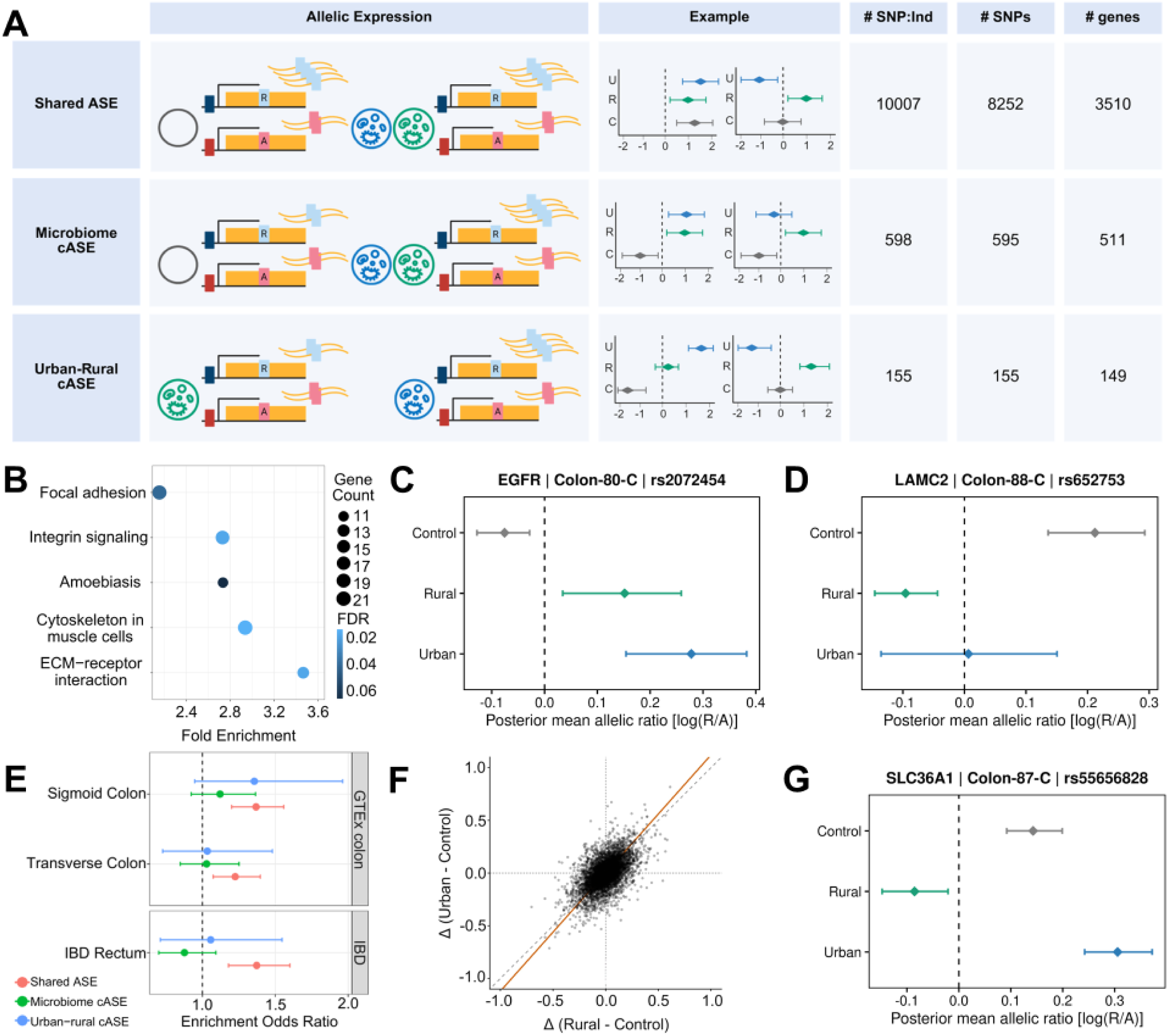
Microbiome exposure modulates host allele-specific expression. **A)** Classification of allele specific expression across microbiome conditions: Allele-specific expression was quantified from reference (R) and alternate (A) allele RNA-seq read counts at heterozygous SNPs within each individual/colonoid line and condition. SNP:individual pairs were then classified as shared ASE, microbiome cASE or urban-rural cASE based on posterior allelic effects across control (C), urban (U) and rural (R) microbiome conditions. Representative posterior means and 95% credible intervals are shown for each class. Numbers indicate the total SNP:individual pairs, unique SNPs and unique genes in each category. **B)** Dotplot of top enriched KEGG pathways for genes exhibiting microbiome-dependent cASE (FDR < 10%). The Y-axis is the KEGG pathway and X-axis is the fold enrichment of genes belonging to the KEGG pathway in microbiome-dependent cASE genes. The size of the dot corresponds to the number of cASE genes associated with the KEGG pathway and the color of the dot indicates the BH p-adjust value for the enrichment test. **C,D)** Examples of loci exhibiting shared microbiome-dependent cASE: X-axis is the posterior mean allelic imbalance effect (indicated by the diamond point) for urban (blue), rural (green) and control (gray) conditions as estimated by mashR. The error bars represent the 95% credible intervals calculated by sampling the posterior distributions for each effect. **E)** Fisher’s exact test of enrichment of either the genes with shared ASE, genes with shared microbiome dependent cASE or urban-rural specific cASE in GTEx Transverse Colon and Sigmoid Colon and IBD Rectum eGene datasets. Error bars indicate 95% confidence intervals. **F)** Scatterplot comparing the difference in posterior mean allelic imbalance between Rural microbiomes and controls (x-axis) and Urban microbiomes and controls (y-axis) across all tested SNP:individual pairs. The dashed gray line indicates the identity line (y = x) and the solid orange line represents the total least squares (TLS) regression fit through the origin. **G)** Example locus exhibiting urban rural cASE: X-axis is the posterior mean allelic imbalance effect (indicated by the diamond point) for urban (blue), rural (green) and control (gray) conditions as estimated by mashR. The error bars represent the 95% credible intervals calculated by sampling the posterior distributions for each effect.

Over-representation analyses revealed significant enrichment for genes involved in epithelial architecture, cell-matrix interactions and structural integrity (**Fig 3B**, FDR < 10%). For example, *EGFR* and *LAMC2* are two host genes that harbor microbiome cASE (**Fig 3C-D**) and both are involved in the focal adhesion and integrin signaling pathways, with *LAMC2* additionally represented in the ECM-receptor interaction and amoebiasis pathways. *EGFR* encodes the epidermal growth factor receptor which activates downstream pathways to promote cell proliferation and migration. *LAMC2* encodes a laminin protein that is needed to anchor epithelial cells to the basement membrane.

Previous studies showed that ASE recapitulated *cis*-regulatory effects detected through eQTL mapping whereas context-dependent ASE were not well captured by studies that do not explicitly model the relevant environmental contexts^30,31^. Analogously, we observed that genes with shared ASE were enriched among genes with eQTLs (eGenes) from three different eGene datasets: the Sigmoid Colon and Transverse Colon datasets in GTEx ^23^, and an eQTL dataset in Rectum tissues from patients with IBD and controls from the Inflammatory Bowel Disease Multi’Omics Database^17,26^(GTEx Sigmoid Colon Fisher’s test OR = 1.36, *P* = 2.0 × 10^−6^; GTEx Transverse Colon Fisher’s test OR = 1.22, *P* = 2.23 × 10^−3^; IBD Rectum eGenes (Fisher’s test OR = 1.37, *P* = 3.18 × 10^−5^; **Fig 3E**). Conversely, neither microbiome cASE nor Urban-Rural cASE were significantly enriched in any eGene dataset. We additionally examined if genes with cASE recapitulated previous findings of host genotype-by-microbiome (GxM) interactions in intestinal biopsies^26^. Of the 27 genes that were previously identified to harbor GxM for specific microbial taxa and that were also tested in this analysis, 3 genes did not exhibit ASE, 20 genes exhibited shared ASE, 3 exhibited microbiome cASE and 1 gene exhibited Urban-Rural cASE. These findings demonstrate that several genes previously implicated in host GxM interactions also exhibit allele-specific effects in the colonoid co-culture system.

To further investigate host allelic effects across the Urban and Rural microbiomes, we focused on the 149 genes that exhibited Urban-Rural cASE, and split them into two subcategories based on the direction of allelic effects relative to control: magnitude cASE, in which Urban and Rural microbiomes altered allelic imbalance in the same direction (90 SNPs in 86 genes), and directional cASE in which Urban and Rural microbiomes altered allelic imbalance in opposite directions (65 SNPs in 63 genes) (**Supplemental Figure 4**), respectively. The greater number of magnitude cASE suggests that lifestyle-associated variation in gut microbiome composition more frequently altered the strength of existing allelic effects rather than their direction. Consistent with this pattern, Urban microbiomes generally induced larger changes in allelic imbalance compared to Rural microbiomes, relative to control (TLS slope 1.12, 95% 1.09 - 1.15, **Fig 3F**). Although Urban-Rural cASE genes were not significantly enriched for Gene Ontology biological processes or KEGG pathways, several genes with established roles in intestinal biology exhibited Urban-Rural cASE, including *SLC36A1*, which encodes a proton-coupled amino acid transporter (**Fig 3G**). Together, these findings demonstrate that compositionally distinct Urban and Rural microbiomes differentially modulate host allelic effects, predominantly through differences in effect magnitude rather than direction.

Shifts in gut microbiome composition associated with industrialization have been linked with increased prevalence of several non-communicable diseases, including obesity, diabetes and inflammatory bowel diseases (IBD)^14^. Transcriptome-wide association studies have identified disease genes by associating their genetically predicted gene expression with disease status in GWAS participants^34,35^. These studies provided a link between genes and diseases mediated by gene expression, and identified putative causal genes by integrating TWAS with colocalization approaches^36^. To investigate whether gene-expression mediated disease risk may be modulated by GxM, we asked whether cASE genes are enriched in disease genes in S-Predixcan. We found that 23 Urban-Rural cASE genes were significantly associated with IBD traits and twelve of these genes (*ACHE, APPL2, ATG16L1, ATG4B, BCR, CHRNB1, EPB41, GNL3, PIEZO1, SDCBP2, STAM2* and *TPM1*) also exhibited significant associations between allelic imbalance and specific microbial taxa (FDR < 5%). Together, these findings demonstrate that lifestyle-associated variation in gut microbiome composition is associated with differences in host *cis*-gene regulation in genes independently associated with IBD, therefore providing a putative mechanism for microbiome disease risk mediated by effects on host gene expression.

### Prevotella copri abundance modulates host genetic effects on gene expression

To identify microbial taxa that may contribute to differential allelic effects between Urban and Rural microbiomes, we leveraged the naturally occurring variation in taxa composition across the eight microbiome samples used in this study. We quantified the correlation between microbial taxon abundance and allelic imbalance for the Urban-Rural cASE loci across the eight microbiome replicates. We identified 2028 significant SNP-taxon associations (FDR < 10%) and focused on the top 549 (FDR < 5%) involving 76 microbial taxa (out of 136 tested) and 83 cASE loci, corresponding to 80 host genes and 546 unique taxon-gene pairs (**Supplemental Figure 5**). Individual cASE loci were frequently associated with multiple microbial taxa, consistent with the strong covariance in microbial composition across the microbial communities. To account for this correlated structure, we clustered the 136 microbial taxa based on the correlation of their abundances across the microbiome samples, identifying 12 taxon clusters (**Supplemental Figure 6**). Seventy-eight of the 83 cASE loci were associated with multiple taxa belonging to at least one microbial cluster. Furthermore, these associations commonly spanned clusters, with 59 loci associated with taxa from two clusters and 16 loci with taxa from three different clusters. Associations were dominated by two strongly anti-correlated groups, a *Prevotella* rich cluster (cluster 2) and a *Bacteroides* rich cluster (cluster 5), with 55 of the 83 cASE loci having significant correlations with taxa from both clusters. Hierarchical clustering of the taxon-cASE correlation profiles similarly revealed opposing patterns of correlations across these microbial groups (**Supplemental Figure 5**). Together, these results suggest that Urban-Rural cASE can be traced back to microbial taxa representative of shifts in microbial community composition characteristic of the rural to urban lifestyle transition.

To independently validate interactions between host genotype and individual taxa, we experimentally tested whether perturbation of a single taxon could recapitulate the observed associations between allelic ratio and microbial abundance. We focused on twelve Urban-Rural cASE loci associated with the abundance of at least one *Prevotella* taxon (FDR < 10%) in the primary experiment. The direction of these associations was concordant across *Prevotella* taxa. We performed a spike-in experiment with *Prevotella copri,* one of the most abundant *Prevotella* species^37^. We supplemented the microbial isolate into a baseline microbial community at 0%, 5% or 50% of the total microbial inoculum and co-cultured with a colonoid monolayer from one donor (n = 3 per condition, **Fig 4A**). Of the 12 Urban-Rural cASE loci, seven exhibited allelic ratio changes concordant with their original *Prevotella* associations at one or both *P. copri* doses (**Supplemental Figure 7**). Four of these Urban-Rural cASE loci showed significant changes in allelic ratio following *P. copri* supplementation: rs34127355 in *ATG4B* (mean allelic ratio difference = 0.34, one-sided t-test *P* = 0.036), rs143156892 in *DYRK1A* (mean allelic ratio difference = 0.35, *P* = 0.013), rs6793735 in *PLS1* (mean allelic ratio difference = 0.12, *P* = 0.031), and rs17355294 in *RUFY1* (mean allelic ratio difference = - 0.31, *P* = 0.036) (**Fig 4B-I**). In the primary experiment, the allelic ratio at all four loci was significantly associated with the abundance of multiple *Prevotella* taxa, including significant associations with *P. copri*. In particular, *ATG4B* exhibited allelic imbalance associated with the abundance of twelve *Prevotella* taxa in the primary experiment. This gene is a core regulator of autophagy and was also significantly associated with IBD by S-Predixcan in both the sigmoid and transverse colon.

**Figure 4.**
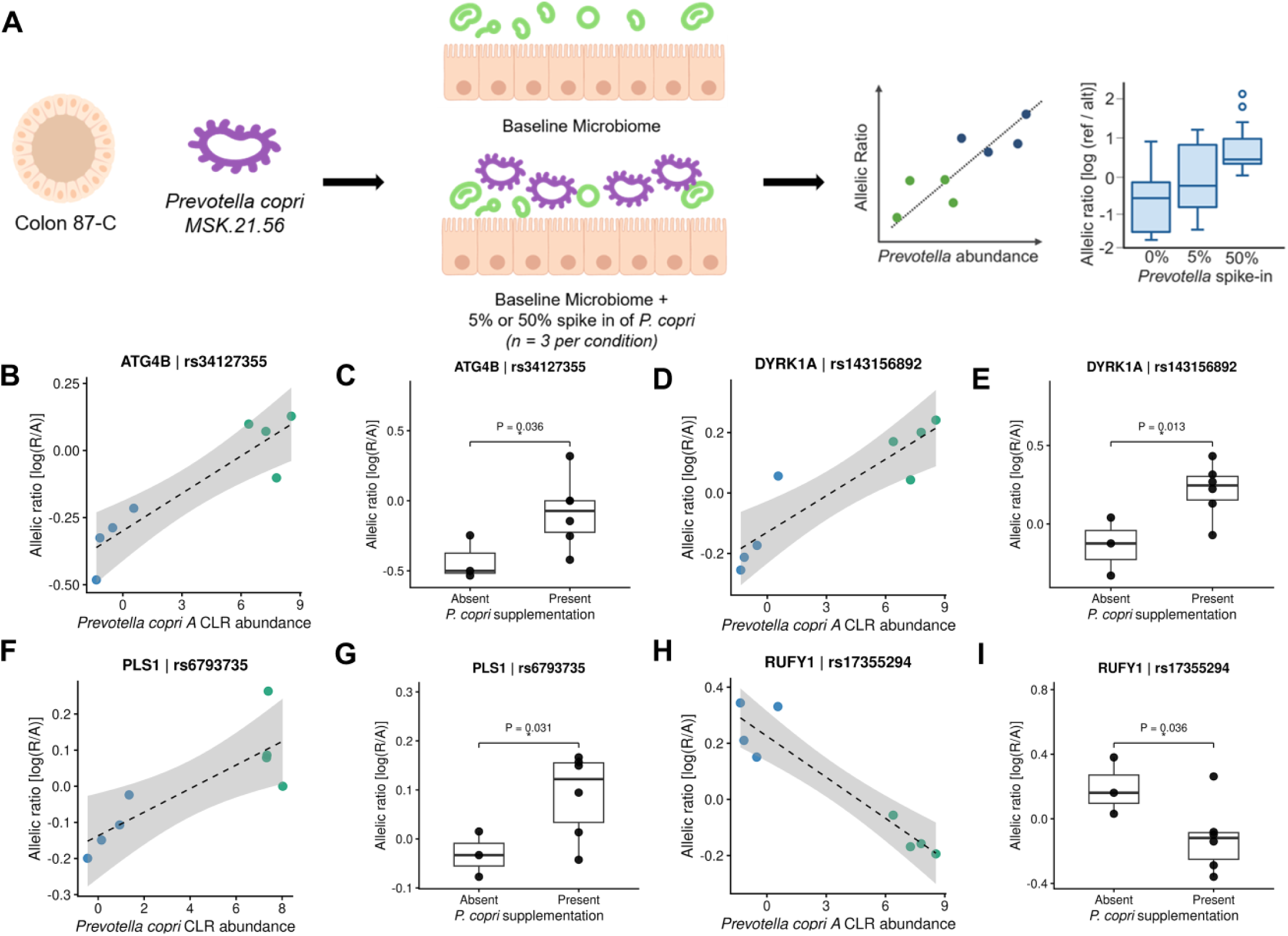
Experimental validation of microbe-induced host allelic expression. **A)** Schematic of the *P.copri* spike-in experiment. Colon-87-C colonoids were treated with a baseline microbial community alone or supplemented with *P.copri* strain MSK.21.56 at 5% or 50% of the total microbial inoculum (n = 3 per condition). Allelic ratio was then quantified following co-culture to test whether experimental increases in *P.copri* abundance recapitulated Prevotella-associated allelic effects observed in the primary experiment. **B, D, F, H)** Representative associations between *Prevotella* abundance and allelic ratio in the primary microbiome experiment for the four loci exhibiting significant effects in the spike in experiment. X-axis is CLR-transformed abundance of microbial taxon and Y-axis is allelic ratio calculated as the natural log of the ratio of reads mapping to REF over ALT allele. Points represent individual microbiome exposure replicates and are colored by urban- or rural-association. Dashed lines indicate linear fits with 95% confidence intervals. **C, E, G, I)** Allelic ratio following *P. copri* MSK.21.56 spike-in for 0% (*P. copri* absent) or 5% and 50% (*P. copri* present) for the four loci exhibiting significant effects in the spike in experiment. Points represent individual experimental replicates (n = 3 for 0% and n = 6 for 5% and 50% together) and boxplots show distribution across replicates. Nominal P value calculated and displayed using one-sided Student’s t-tests, with the direction specified by the original *Prevotella*-allelic ratio association. P < 0.05; ns, not significant.

## Discussion

Differences in the composition of the gut microbiome have been observed across different lifestyles^5–11^ and disease states^3,4,15^. Previous studies aimed to characterize the functional consequences of these differences through host transcriptional analyses^16–22^. However, as genetically distinct individuals transition through lifestyles, microbiome compositional differences may impact gene regulation in a host genotype-dependent manner. Here, we used a co-culture system with a panel of colonoids from healthy donors and live microbial communities representative of urban and rural lifestyles to characterize both host average transcriptional and allele-specific responses to microbiome variation. Urban and Rural microbiomes elicited largely shared transcriptional responses, although urban microbiomes generally induced larger changes in gene expression. In contrast, allele-specific expression analyses revealed substantial heterogeneity in *cis*-regulatory genetic effects with hundreds of loci exhibiting microbiome-dependent allele-specific expression and a subset of variants with differential allelic responses to urban and rural microbiomes. These allelic effects were associated with differences in microbial community composition, and experimental perturbation of *Prevotella copri* recapitulated allele-specific expression effects associated with this taxa abundance in the Urban and Rural microbial communities. Together, these findings demonstrate that microbiome composition can modulate host gene regulation in a host genotype-dependent manner and illustrates the utility of experimentally tractable systems and allele-specific expression approaches for identifying host-microbiome interactions.

An important feature of our experimental system is its ability to capture biologically relevant intestinal epithelial responses to live microbial communities. The host transcriptional response to microbiome exposure was characterized primarily by downregulation of host antiviral and type I interferon signaling, including canonical interferon-stimulated genes, as well as metallothionein-associated responses. These pathways are consistent with established functions of the intestinal epithelium, which regulates innate immune signaling to microbial exposure^38^, while metallothioneins contribute to host immune regulation and protection from cellular stress^39^. More broadly, bidirectional crosstalk between the gut microbiome and intestinal epithelium contributes to epithelial function and homeostasis^40^, and disruption of these interactions has been implicated in different pathologies including pathogen infections ^41^ and Inflammatory Bowel Disease^42^. Intestinal organoids, including colonoid-derived monolayers, have emerged as tractable models for studying host-microbe interactions^43^, including epithelial responses to invasive enteric pathogens^44^, and transcriptional responses to complex microbial communities in the setting of Ulcerative Colitis^45^. Our findings extend this framework by examining the role of host *cis*-regulatory genetic variation in epithelial responses to microbial exposure. In particular, genes that harbored microbiome cASE were enriched in pathways involved in epithelial architecture and structure, suggesting that microbiome-epithelial crosstalk can vary with host genetic background. These findings introduce host *cis* gene regulation as an additional dimension of host epithelium-microbial cross talk and demonstrate the utility of our colonoid co-culture model as a meaningful system for investigating genotype-by-microbiome effects.

Allele-specific expression (ASE) provides a means to uncover host *cis*-regulatory genetic effects on gene expression that dynamically vary across microbial community contexts. While genes exhibiting ASE across control, Urban and Rural microbiome conditions were enriched among eGenes identified in GTEx colon and IBD Rectum datasets, genes that harbored microbiome cASE and urban-rural cASE were not. This is consistent with previous work demonstrating that explicitly comparing ASE across multiple environments can reveal *cis*-regulatory genetic effects on gene expression that are missed when genetic effects are examined in an arbitrary “baseline” context^30^. When considering the gut microbiome, such effects are particularly relevant as individuals are continually exposed to microbial communities that vary across lifestyle, environment and over time, thus potentially inducing changes in genetically-regulated host gene expression. Our results demonstrate that a large number of genes are under the joint regulatory control of host genetics and the microbiome and highlight the importance of measuring and modeling the microbial context when characterizing host *cis*-regulatory genetic variation.

Lifestyle-associated differences in microbiome composition also influenced the magnitude of host allelic effects. Urban-Rural cASE loci more frequently reflected differences in the magnitude rather than the direction of allelic effects, with Urban microbiomes generally eliciting larger changes in allelic imbalance relative to control. Urbanization is accompanied by substantial restructuring of the gut microbiome, including depletion of *Prevotella* species with greater representation of Bacteroides and other taxa^6–10,22,32^. Because differences in microbial community composition can alter the microbial signals and metabolites encountered by the host intestinal epithelium^40^, the greater host transcriptional and allelic responses elicited by Urban microbiomes may reflect broader functional consequences of urbanization-associated shifts in microbiome composition on host epithelial homeostasis. Importantly, the differential allelic effects observed here further suggest that the magnitude of host responses vary with host genetic background. Thus, lifestyle-associated variation in gut microbiome composition may influence not only the overall strength of host intestinal epithelial responses but also interindividual variation in those responses through interactions with host genotype.

The association of Urban-Rural cASE with microbial community composition provides further evidence that differences in host genetic effects are linked to lifestyle-associated microbiome variation. Associations between allelic imbalance and microbial abundance were dominated by strong anti-correlations between *Prevotella-* and *Bacteroides*-rich groups of taxa. Individual cASE loci were frequently associated with multiple taxa, especially within and between the *Prevotella* and *Bacteroides* clusters, reflecting the extensive co-abundance within these microbial communities and making it difficult to attribute allelic effects to individual taxa. However, targeted perturbation of *Prevotella copri* allowed us to test whether a single member of the correlated *Prevotella* community could contribute to the allelic effects observed. Notably, all four loci significantly associated with *P. copri A* abundance showed directionally concordant responses to *P. copri* supplementation, with three reaching statistical significance. One additional locus associated with *P. copri* but not specifically *P. copri A*, was significantly recapitulated, while another *P. copri* - associated locus did not exhibit a concordant change in allelic ratio. In contrast, none of the six loci that lacked associations with *P. copri or P. copri A* were significantly recapitulated. This pattern suggests that the experimental effects were preferentially recapitulated at loci whose allelic expression correlated with *P. copri* abundance. Furthermore, these results suggest that these regulatory effects may depend on functional differences among *Prevotella* strains, interactions among multiple taxa or the broader microbial community context. These results demonstrate that a joint association-based and spike-in experimental approach can characterize specific taxa interacting with host genotype, confirming that individual microbial taxa can contribute to host genotype-dependent effects on gene regulation, and paving the way for future microbe-specific characterization of host-microbiome interactions in gene regulation.

The convergence between cASE associated with lifestyle-variation in microbiome composition and genes implicated in IBD provides potential disease relevance for these context-dependent genetic effects. Urbanism-related shifts in gut microbiome composition have been previously linked to non-communicable disease prevalence through epidemiological studies^46–48^, while host genetic variation has also been associated with risk for many of the same diseases^49–51^. The overlap of Urban-Rural cASE with IBD-associated genes suggests that *cis*-regulatory genetic effects responsive to lifestyle-associated variation in microbiome composition can occur in genes independently implicated in intestinal disease. Notably, *ATG4B*, is relevant in this context as a core regulator of autophagy, as it has an established role in intestinal homeostasis and has been found to be protective against experimental colitis^52^. It exhibited cASE associated with multiple *Prevotella* species including *Prevotella copri A*, a species previously implicated in IBD^53^. The experimental validation of this association with *Prevotella copri* further supports a specific microbial contribution to the observed allelic response. These complementary associations highlight how microbiome composition can shape host genetic effects within biological processes relevant to intestinal disease.

While this study establishes a controlled experimental framework to investigate microbiome-dependent host gene regulation, there are limitations to the resolution of these host-microbiome interactions that can be detected. The relatively small number of donor-derived colonoid lines and microbial communities capture only a subset of host genetic and lifestyle-associated microbiome variation. Strong covariance in composition across the microbial communities limits the ability to attribute effects to individual taxa. The *P. copri* spike-in experiment begins to address this limitation through targeted perturbation but was performed in a single colonoid line with a limited number of replicates and candidate loci. Consequently, the significant responses observed provide experimental support for specific candidate effects rather than broad validation of all *Prevotella*-associated cASE. Furthermore, although epithelial cells are the primary interface between the host and gut microbiota, the colonoid system does not capture interactions with immune, stromal and other cell types that may contribute to host responses. Future studies incorporating larger panels of host genotypes and microbial communities, targeted perturbations with other defined microbial taxa, and more complex co-culture or *in vivo* systems will therefore be important for further resolving the host and microbial factors underlying genotype-dependent regulatory effects.

In conclusion, our findings demonstrate that gut microbial communities can modify host *cis*-regulatory genetic effects on gene expression in human intestinal organoids and that these effects vary with lifestyle-mediated shifts in microbiome composition. Combining allele-specific expression with an experimentally tractable co-culture system with colonoids and live microbial communities enabled the detection of microbiome-dependent host genetic effects that are associated with microbiome community composition and can be tested through targeted perturbation with specific microbial taxa. Expanding this framework across greater host genetic and microbial diversity will help to unravel the mechanisms underlying genotype-by-microbiome interactions and their contributions to human phenotypes and disease.

## Methods

### 3D Colonoid Culture

Experiments were conducted using four primary human colonic organoids (colonoids) isolated from healthy adult donors (age 21-59) obtained from MilliporeSigma (MilliporeSigma Cat# SCC312, SCC321, SCC323, SCC330) (**Table 1**). All lines were reported by the vendor to be mycoplasma negative, from normal healthy tissue and negative for infectious diseases prior to shipment.

**Table 1.** Colonoid Line Information.

| Catalog Number | Line | Sex |
|---|---|---|
| SCC312 | Colon-80-C | F |
| SCC321 | Colon-87-C | M |
| SCC323 | Colon-88-C | F |
| SCC330 | HT-099-C | F |

Colonoid culture and differentiation protocols were adapted from previously published methods developed by the University of Michigan Human Organoid Core Facility (RRID:SCR_027333)^54,55^. Three-dimensional (3D) colonoid cultures were maintained in Cultrex UltiMatrix growth factor reduced basement membrane extract (R&D Systems, BME001-05), diluted to final concentration of 2.5 mg/mL in culture media, embedded as 10 µL domes in 12-well or 6-well plates (ThermoFisher,140675 and 150628). Domes were polymerized at 37⁰ C for 10 minutes prior to overlay with media. To promote a cystic organoid phenotype, colonoids were cultured in IntestiCult Organoid Growth Medium (StemCell Technologies, 06010) supplemented with 2.5 µM CHIR99021 (Tocris-Fisher, 4423). Additionally, media was supplemented with 10 µM Y-27632 (ROCK Inhibitor) (Tocris, 125410) and 100 µg/ml Primocin (Invivogen, ant-pm-1). Colonoids were cultured at 37°C and 5% CO_2._ Media was changed daily and colonoids were passaged every 5-7 days at approximately 1:1.5 to 1:2 split ratios by mechanical disruption.

To generate sufficient cells for downstream monolayer differentiation, colonoid lines were passaged and expanded in 3D culture until approximately 4 wells of a 6-well plate contained confluent colonoid domes. Experiments were performed using colonoids between passages 8 and 21.

### Colonoid 2D Monolayer Differentiation

Two-dimensional (2D) monolayer differentiation was performed using protocols adapted from previously published methods developed by the University of Michigan Human Organoid Core Facility (RRID:SCR_027333) ^54,55^. Briefly, 3D colonoids were harvested and washed with ice-cold 0.5 mM EDTA in DPBS, and dissociated with 0.05% Trypsin-EDTA (1X) (Gibco 25300-054) at 37°C for 3.75 minutes. After inactivation of trypsin with 10% FBS in Advanced DMEM/F-12, cells were mechanically dissociated by repeated pipetting and then passed through a 40-µm nylon filter to produce a single-cell suspension and remove large aggregates. Cells were seeded into 96-well plates (ThermoFisher 1256566), coated with collagen IV at 10µg/cm^2^ (Sigma C5533), at 400,000 cell aggregates per well in IntestiCult Organoid growth medium supplemented with 10 µM Y-27632 (ROCK Inhibitor) (Tocris, 125410) and 100 µg/ml Primocin (Invivogen, ant-pm-1). Monolayers were cultured at 37°C and 5% CO_2_ for 24 hours prior to differentiation. Following attachment after 24 hours (Day 0), the 3D growth medium was replaced with 2D differentiation media which contained Advanced DMEM/F-12 (FisherScientific 12634028), 4 mM GlutaMAX (Gibco 35050061), 20 mM HEPES buffer (FisherScientific 15630080), 2X N-2 media supplement (FisherScientific 17502048), 2X B-27 Supplement Minus Vitamin A (FisherScientific 12587010), 2mM N-Acetyl-L-cysteine (Sigma A9165), 50ng/mL human EGF (R&D 236-EG-01M), 50 ng/mL Noggin (R&D 6057-NG-025/CF), 10 nM Gastrin (Sigma G9020), 2.5 µM Y-27632 and 50 units/mL Penicillin/Streptomycin (Gibco 15140122). On Day 2, monolayers were switched to complete differentiation medium without Y-27632 and Penicillin/Streptomycin, on day 3 (24 hours prior to the co-culture experiment) they were moved to a low oxygen incubator (5% O_2_, 5% CO_2_, 37°C). Monolayer morphology and confluency were monitored by brightfield microscopy each day.

### Live Microbiome Sample Selection

Fecal sample processing, microbiome extraction, storage, metagenomic sequencing and OTU table generation have been previously described^22,32^. Briefly, fecal samples from healthy adults across four countries (Ghana, Rwanda, Nigeria and Malaysia) were collected as part of the Global Microbiome Conservancy (GMbC) initiative (http://microbiomeconservancy.org) and processed as previously described, with whole live microbial communities extracted from the samples under anaerobic conditions and stored in glycerol at -80°C until use.

For this study, a subset of eight samples from this cohort were selected for colonoid co-culture experiments (see co-culture methods below). To minimize potential confounding with geographic and cultural variation, samples were selected from donors living in Ghana and Rwanda, with two Urban and two Rural microbiomes from each country. Urban and rural classifications were assigned using previously described population density thresholds^11^, where samples collected from participants in communities with densities > 1,000 inhabitants / km^2^ were classified as urban.

Rural samples were from donors maintaining traditional farming lifestyles while urban samples were from donors who transitioned from farming to westernized urban lifestyles. This study design allowed for within-country urban/rural contrasts while reducing broader geographic confounding and enriching for microbiome variation associated with urbanization-related lifestyle transitions. Full cohort characteristics for the 8 individuals can be found in supplementary table 1.

### Microbiome Profiling and Analysis

#### Taxon filtering and transformation

Species-level metagenomic abundances from the full microbiome sample cohort were filtered to retain taxa present at ≥0.001 relative abundance in at least 25% of Urban or Rural samples and transformed using the centered log-ratio (CLR) as done in Arif et al 2025^22^. This yielded 136 microbial taxa. CLR values for the eight microbiome samples used in this study were then extracted for all downstream analyses.

#### Community composition

Microbiome community composition was compared across the eight samples at the species level using principal coordinates analysis (PCoA). Microbial abundances from the OTU table (136 taxa total) were CLR-transformed and Euclidean distance was used to compute a sample-wise distance matrix. PCoA was performed on this distance matrix using the pcoa() function from the ape package.

#### Differential abundance of taxa associated with urbanization

To evaluate whether taxa previously reported to differ between Urban and Rural populations also differed in our samples, the relative abundances of all species belonging to the genera *Bacteroides* and *Prevotella* and the phylum Firmicutes were summed for each sample. Differences between the four Urban and four Rural microbiomes were assessed using a one-sided Wilcoxon rank-sum test, with the direction of the alternative hypothesis specified from prior studies^6–11,22,32^. P < 0.05 was considered significant.

### Experimental Design and Replicates

Colonoid lines were differentiated in two experimental batches based on line growth and amount of organoids. Each experimental batch contained two colonoid lines processed in parallel. Each colonoid line was seeded into an independent 96-well plate and maintained as a separate experimental unit within each batch. Within an experimental batch, all colonoid lines were differentiated, co-cultured with the microbiome samples, processed for RNA extraction, prepared for library construction and sequenced together.

For each colonoid line, 2 wells representing technical replicates were seeded for each of the 12 conditions consisting of four urban-associated microbiomes, four rural-associated microbiomes and four media-only control replicates, for a total of 24 wells. Technical replicates were pooled prior to library preparation yielding one RNA-sequencing library per colonoid line and treatment condition.

### Colonoid-Microbiome co-culture

We adapted our published colonocytes-microbiome co-culture protocol^19–22^ to perform colonoid monolayer-microbiome co-culture experiments. The 8 live microbial communities extracted from the fecal samples were thawed on ice for 30 minutes and then quickly at 37°C. The microbial samples were diluted 1:2 in PBS (Gibco 10010-023) and then the microbial density was assessed using the OD600 measured with a spectrophotometer (Bio-Rad SmartSpec 3000). Each microbial sample was then diluted in Complete Differentiation Media without antibiotics in order to apically treat the colonoid monolayers at a 1:10 ratio of host cells to microbes for 2 hours at 37°C and 5% O_2_. Control wells were incubated with media with no microbiome in the same conditions in parallel. Following incubation, plates were placed on ice, wells were rinsed 2X with cold PBS and then cells were lysed within each well using Lysis/Binding Buffer (Invitrogen) and frozen at -80°C until library preparation.

### Host RNA sequencing and processing

For RNA library preparation, plates within an experimental batch were processed in parallel. Poly-adenylated mRNAs were isolated from thawed lysates using the Dynabeads mRNA Direct Kit (Invitrogen 61012) and following manufacturer instructions. RNA-seq libraries were prepared using a protocol modified from the NEBNext Ultra II Directional RNA Library Preparation Kit (NEB E7760L) with sample-specific barcodes from BIOO Scientific that were added by ligation^56^. Libraries generated within each experimental batch were pooled and quantified using both the Agilent Bioanalyzer and Qubit fluorometric quantification (ThermoFisher Scientific). Libraries were pooled and sequenced on an Illumina NovaSeq 6000 at the Genome Sciences Core at Wayne State University using an S2 flow cell with 300 cycles to generate 2 x 150 bp paired-end reads. Sequencing was performed with a target depth of 100 million reads per library. The number of reads per library can be found in Supplemental Table 2.

HISAT2^57^ was used to align demultiplexed RNA-seq reads to the human genome version ‘GRCh38_snp_tran’ which considered splicing and common genetic variants. Samtools (version 1.19)^58^ was then used to generate clean bam files by removing PCR duplicates and reads with a quality score of <10 (equating to reads mapped to multiple locations).

### Differential gene expression analysis

#### Preprocessing

Gene counts were generated with HTSeq^59^ and the Ensembl release 103 GRCh38 transcriptome assembly across 60612 genes. Counts were obtained from position-sorted BAM files using reverse-stranded counting (--stranded == reverse). Gene count matrices were filtered to retain autosomal protein-coding genes with non-zero expression in at least 50% of all samples included in a given analysis.

#### DESeq2 Analysis

Differential gene expression analyses were performed independently within each colonoid line using DESeq2 (v1.32.0)^60^ in R(v4.3.2) and comparing microbiome-treated samples to control using the design formula *Expression* ∼ *Treatment*. Initial analyses revealed substantial heterogeneity in microbiome-induced transcriptional response across donor lines, ranging between 57 and 3054 differentially expressed genes (FDR < 10%, DESeq2, Supplemental Figure 1). In order to identify the transcriptional response to microbiome exposure that is shared across different hosts, we combined log2 fold change estimates and standard errors from DESeq2 for each of the four lines using an inverse-variance weighted fixed-effect meta-analysis.

To identify transcriptional responses specific to Urban or Rural microbiome exposures, i.e. Urban microbiomes versus control and Rural microbiomes versus control, data was subsetted to the relevant replicates (Urban or Rural microbiome-treated replicates, respectively, and control replicates) and DESeq2 was run using the model *Expression* ∼ *Treatment* within each line followed by meta-analysis. Importantly, no shrinkage step (“lfcShrink”) was performed because of the subsequent meta-analysis and then re-analysis using MASH (see below). Additionally, we directly contrasted Urban versus Rural microbiome responses, by subsetting to Urban or Rural microbiome-treated replicates within each line, and identifying differential gene expression using the design formula *Expression* ∼ *Urbanism* followed by meta-analysis. For all these analyses, 16,000 genes were tested across all colonoid lines. Multiple testing correction was performed using Storey’s q-value and genes with a q-value < 0.10 were considered significantly differentially expressed. Full DESeq2 results can be found in Supplemental Tables 3 - 22.

#### MASH Analysis of differential gene expression

We used MASH (mashR R package version 0.2.79^33^) to jointly model differential gene expression between urban microbiomes and control, and rural microbiome and controls, respectively to better understand the relationship in transcriptional responses between the two lifestyle associated microbiome exposures. Specifically, we analyzed the log-fold changes and corresponding standard errors for 16,000 genes from the meta-analyses above. Our MASH analysis closely followed the recommended analysis pipeline for re-analyzing eQTL results in multiple conditions (https://stephenslab.github.io/mashr/articles/eQTL_outline.html). Importantly, because the LFCs are estimated against a common control condition, it was important to account for the strong “null” correlation structure among the urban and rural conditions; this correlation was estimated to be 0.52. MASH re-estimates, or “shrinks”, the LFCs by generating posterior estimates of the LFCs, similar to the “shrinkage” performed in other differential expression analyses, the key difference being that MASH shrinks the LFC estimates jointly in the two conditions according to sharing patterns learned from the data (this includes a “null” sharing pattern). MASH learned that only three covariance patterns were needed to explain the pattern of LFCs in the urban and rural conditions (weight > 0.1%): an “equal effects” covariance in which the LFCs are the same; a second covariance pattern in which the LFCs in the urban condition were slightly larger; and a “null” covariance in which all LFCs were zero. Posterior effect size estimates and significance (lfsr) output from MASH can be found in Supplemental Table 23.

#### Visualization

For visualization of selected differentially expressed genes, raw count data were transformed using the variance stabilizing transformation (VST) in DESeq2. To facilitate comparison of expression across donors, transformed expression values were adjusted using the removeBatchEffect() function from the limma package^61^ with colonoid line (donor) specified as the batch variable. Batch-corrected expression values were used exclusively for visualization of gene expression patterns, while all differential gene expression analyses were performed on the original count data.

### Over-representation analyses of differentially expressed host genes

To determine host biological processes and pathways that were over-represented in significantly differentially expressed genes, results were first separated into upregulated and downregulated genes based on the log2FoldChange sign. The ClusterProfiler^62^ R package was then used to perform biological process Gene Ontology^63,64^ over-representation analyses with the background gene universe defined as all tested genes. Biological processes with FDR < 10% were defined as significantly enriched. Full enrichment results can be found in Supplemental Tables 24 - 29.

### Genotyping and Imputation

Genomic DNA was isolated from colonoid lines using the DNeasy Blood &Tissue Kit (Qiagen, 69504) according to the manufacturer’s instructions. DNA samples were submitted to Genewiz (Azenta Life Sciences) for low-coverage (∼1X) whole-genome sequencing. Genotypes were subsequently imputed using a GLIMPSE2-based imputation pipeline^65^ against the 1000 Genomes Project reference panel^66^. Imputed genotype calls were converted to posterior genotype probabilities and used for downstream ASE analyses.

### Allele-Specific Expression Quantification

Allele-specific expression (ASE) was quantified at heterozygous biallelic SNPs for each donor. Heterozygous SNPs were identified from imputed genotype data by retaining variants with a genotype probability (GP) > 0.999 filtered for common SNPs from dbSNP build 144. RNA-seq alignments from all samples belonging to a given donor were jointly processed using bcftools mpileup and bcftools call (bcftools v1.19) to generate allele-specific pileups across all samples from that donor. Joint processing across samples was used to improve variant detection and ensure consistent SNP representation across conditions. Resulting pileups were filtered to retain only biallelic SNPs and then converted to a long-format allele count table containing one row per SNP-sample pair with reference and alternate allele counts for all SNP-samples across all donors. SNP-sample pairs were first filtered to retain those with total read coverage > 40 reads with at least one read mapping to the reference and alternate alleles. To ensure sufficient number of replicates in each condition, analyses were restricted to SNP-donor pairs that passed the filter for at least two control samples, two urban microbiome samples and two rural microbiome samples.

For each remaining SNP-donor pair, allelic ratio was quantified within each sample as the natural logarithm of the ratio of reference to alternate allele reads. Within each condition (control, Urban microbiome, and Rural microbiome), allelic imbalance estimates were averaged across replicates to derive the mean allelic imbalance. Standard errors were estimated as the sample standard deviation divided by the square root of the number of replicate samples within a condition. This approach is based on the assumptions for quantifying allele-specific expression as done in Findley et al^30^. SNP-donor pairs exhibiting no variation in allelic imbalance across replicates were excluded leaving 19,715 SNP-donor pairs including 14,302 unique SNPs across 4,845 genes tested. This table was used for downstream ASE and conditional ASE analyses and can be found in Supplemental Table 30.

### Shared and microbiome-dependent ASE inference using MASH

To identify ASE patterns shared across conditions and microbiome-dependent ASE (cASE), we used multivariate adaptive shrinkage (MASH)^33^ to model allelic imbalance across control, Urban and Rural microbiome conditions, respectively. MASH enables joint re-estimation of effect size across multiple conditions, improving effect size estimation and statistical power relative to individual condition-by-condition analyses.

Condition-specific allelic ratio estimates and standard errors were used as input into MASH. The MASH model was fit following the standard MASH workflow using a random subset of 5,000 SNP-individual pairs to estimate the null correlation structure and fit the mixture model. Data-driven and canonical covariance matrices were included to model patterns of sharing across conditions. This model was subsequently applied to all 19,715 tested SNP-individual pairs to obtain posterior estimates.

Posterior distributions, posterior mean effect sizes and lfsr were estimated for each SNP-individual pair in each of the three conditions. A SNP-individual pair was considered to exhibit ASE within a condition if the absolute value of the posterior mean was greater than 0.1 and the lfsr was less than 0.05. Then for the union of SNP-individual pairs with significant ASE in at least one of the three conditions, 1,000 draws were obtained from the posterior distributions of theeffect size estimated by MASH, in each condition separately. These posterior samples were subsequently used to calculate 95% highest posterior density (HPD) intervals by identifying the narrowest interval containing 95% of the posterior probability mass. Overlap or non-overlap of condition-specific HPD intervals were then used to classify SNP-individual pairs into shared and microbiome-dependent ASE categories (**Supplemental Figure 4**). Shared ASE was defined as overlapping 95% HPD intervals between both Urban and control and Rural and control conditions, indicating consistent allelic imbalance across all conditions. Microbiome-dependent cASE was defined as non-overlapping HPD intervals between at least one microbiome condition and the control. From this subset, we further defined Microbiome cASE when Urban and Rural HPD intervals overlapped, indicating a shared microbiome-associated shift in allelic imbalance relative control. Urban-rural cASE was defined as SNP-donor pairs exhibiting non-overlapping urban and rural HPD intervals, and non-overlapping HPD intervals between at least one microbiome condition and control. These were then further defined as Urban-Rural magnitude cASE when Urban and Rural effects relative to control were concordant or Urban-Rural directional cASE when Urban and Rural effects relative to control were discordant. Concordance was defined as *sign*(*pm_U_* − *pm_C_* ) = *sign*(*pm_R_* − *pm_C_* ) where *pm_U_* , *pm_R_* , and *pm_C_* denote the posterior mean allelic imbalance estimates for Urban microbiome, Rural microbiome and control conditions, respectively. The full table with all SNP-donor pairs including posterior means for each condition, significance and classification can be found in Supplemental Table 31.

### ASE and cASE enrichment in eGenes and Pathway Enrichment

We tested for enrichment of genes with shared ASE and microbiome cASE, respectively, in eGenes from the GTEx (v8) Sigmoid Colon and Transverse Colon datasets^23^ as well as eGenes from the Rectum previously identified in patients with IBD from the IBDMDB database^26^. We restricted our analysis to include genes that were tested in both the eQTL study and for ASE and cASE in our study and used Fisher’s exact test to test for enrichment for each comparison.

We also tested for enrichment of shared ASE, microbiome cASE and urban-rural cASE gene sets in different biological processes and pathways. The ClusterProfiler^62^ R package was used to perform biological process Gene Ontology^63,64^ and KEGG^67^ over-representation analyses for each gene set separately with the background gene universe defined as all genes that had at least one SNP tested for ASE. Biological processes or KEGG pathways with FDR < 10% were defined as significantly enriched. Full enrichment results can be found in Supplemental Tables 32 and 33.

### Association of microbial abundance with urban-rural cASE

To identify microbial taxa associated with differences in host gene regulation between Urban and Rural microbiomes, we tested whether allelic imbalance at urban-rural cASE loci covaried with the abundance of the 136 CLR-transformed microbial taxa across the eight microbiomes Allelic ratios were quantified as log(Ref/Alt), where Ref and Alt are the read counts for the reference and alternate allele derived from the pileup pipeline. For loci with allelic ratio measurements across all 8 microbiome treatments (reads > 40) (n = 87 SNP-colonoid line pairs), Spearman correlations were calculated between allelic ratio and CLR abundance of each taxon across microbiome conditions, for a total of 11,832 tests. P-values were adjusted for multiple testing using Benjamini-Hochberg’s p-adjust and significant associations were defined as FDR < 5%. Full results of the correlation analysis can be found in Supplemental Table 34.

To assess whether significant associations were shared among co-abundant taxa, the 136 taxa were clustered based on pairwise Spearman correlations across the 8 microbiome samples used in this study using hierarchical clustering (average linkage, selected based on cophenetic correlation; distance = 1 − ρ). Cluster number (k = 12) was determined using silhouette width and inspection of within-cluster correlation structure (**Supplemental Figure 4**). For each SNP-colonoid pair, we then counted the taxa with significant correlations falling in each cluster.

### *Prevotella copri* spike-in experiment

To experimentally validate that *Prevotella copri* is a main driver of the taxon-associated allelic effects observed in the primary experiment, we leveraged an independent experiment in which a *P. copri* strain MSK.21.56 was supplemented into a baseline microbiome at 5% or 50% of the total microbial inoculum and co-cultured with colonoid line Colon-87-C. A microbial community originally obtained from OpenBiome from a healthy donor (02-028-C, FMB028) was used as the baseline microbiome as it contained a low endogenous relative abundance of *P. copri* (0.0131%). *Prevotella copri* strain MSK.21.56 was obtained from the Symbiotic Bacterial Strain Bank at the Duchossois Family Institute (DFI) in the University of Chicago.

Colon-87-C was expanded in 3D culture and differentiated as 2D monolayers as described above. Experiments were performed at passage 20. Prior to co-culture, sterile PBS and complete differentiation medium without antibiotics were pre-reduced overnight under anaerobic conditions. The baseline microbiome and *P. copri* were thawed quickly under anaerobic conditions and then diluted separately in reduced PBS. Microbial density was estimated by OD600 as described above. The *P. copri* was spiked into the baseline microbial community at 0, 5 and 50% spike-in concentrations. Colonoid monolayers were treated at a 1:10 ratio of cells to microbes in media for 2 hours at 37°C and 5% O. Each treatment was performed in triplicate (3 biological replicates) with two technical replicates seeded per replicate for a total of 18 wells of a 96-well plate. Following treatment, the plate was placed on ice, wells were rinsed 2X with cold PBS and then cells were lysed within each well using Lysis/Binding Buffer and frozen at -80°C until RNA extraction and library preparation were performed, as described above. Libraries were pooled and sequenced on an Illumina NovaSeq X at the Genomics Facility at the University of Chicago on a 10B flowcell with 300 cycles to generate 2 x 150 bp paired-end reads. Sequencing was performed with a target depth of 100 million reads per library (Supplemental Table 35). RNA seq read processing, alignment, and allele specific expression quantification were performed as described above.

### Analysis of allele-specific effects following *Prevotella copri* spike-in

We considered the 12 Urban-Rural cASE loci in Colon-87-C whose allelic ratio significantly correlated with the abundance of at least one *Prevotella* taxon in the primary analysis (FDR < 10%) and with sufficient allele-specific read coverage across all three replicates of each spike-in condition (total reads > 15 per replicate). Allelic ratio was quantified within each sample as the natural logarithm of the ratio of reference to alternate allele reads, consistent with the earlier analyses. We compared the allelic ratio for the spike-in condition (*Prevotella copri* presence at 5% or 50%) to the baseline microbiome condition (*Prevotella copri absence, 0%*), using one-sided Student’s t-tests assuming equal variance, with the direction of each test specified by the direction of the original *Prevotella*-allelic imbalance associations. We additionally tested for dose-specific effects performing the same t-test described above for 0% versus 5%, and 0% versus 50%, as well as tested for dose-dependent effects using linear regression with *P.copri* spike-in proportion (0, 0.05 and 0.50) as a continuous predictor. Note that for loci associated with multiple *Prevotella* taxa, all significant associations were concordant in direction of effect. Spike-in effects were considered concordant when the direction of allelic imbalance difference matched the direction of the corresponding correlations in the primary experiment. Nominal *P* < 0.05 was used to assess evidence of replication. Results for the t-tests and linear model for each SNP can be found in supplemental table 36.

### Overlap with disease genes in S-PrediXcan

To evaluate whether genes exhibiting context-dependent allelic-specific expression (cASE) were associated with complex traits through their predicted expression in intestinal tissue, we considered S-PrediXcan associations for 8 traits spanning IBD subtypes (Inflammatory Bowel Disease, Crohn’s Disease, Ulcerative Colitis) and metabolic phenotypes (fasting glucose, fasting insulin, LDL cholesterol, IDL triglycerides, CH2:DB ratio) from the supplementary data of ^34^. Within each trait-tissue combination, genes were designated as significant S-PrediXcan associations at a Bonferroni-corrected FDR < 5%, accounting for the number of genes tested per tissue. ASE and cASE genes that overlap with S-Predixcan-associated genes for each trait can be found in supplemental table 37.

## Data Availability

Sequencing files have been uploaded to the Sequence Read Archive (SRA) and are being processed, Bioproject ID is pending.

Code is available at: https://github.com/piquelab/GMCcol

## Author Contributions

SN aided in developing the experimental design, selected microbiome samples, led and conducted the experiments, analyzed the data, interpreted the results, prepared the figures, wrote the first draft of the manuscript and revised the manuscript. SA analyzed microbiome data, interpreted the results, provided feedback on the first draft of the manuscript. GG assisted with colonoid culture and experiments, performed microbial culture for the spike-in experiments and performed library preparation for the validation experiments. AA assisted with colonoid culture and experiments, and performed library preparation for the primary experiments. KH aided in developing the experimental design and assisted with colonoid culture for the validation experiments. MP and MG provided the expertise on the microbiome cohort, interpreted the results and provided feedback on the manuscript. PC provided statistical guidance, contributed to the development and implementation of the MASH analysis and provided feedback on the manuscript. RPR provided guidance and supervision on statistical analyses, interpreted the results, provided feedback on the manuscript, and revised the manuscript. RB contributed to the design of the project, supervised the project, interpreted the results, and provided feedback on the manuscript and revised the manuscript. FL designed the project, supervised the project, interpreted the results, provided feedback on the manuscript, revised the manuscript. All authors have read and approved the final manuscript.

## Acknowledgments

We thank all human participants that agreed to provide samples to the GMbC project. We thank Sonia Kupfer and members of the Luca, Pique-Regi, Blekhman, and Kupfer groups for helpful discussions and feedback on earlier versions of this manuscript. We are grateful to the University of Michigan Human Organoid Core and the University of Chicago Center for Interdisciplinary Study of Inflammatory Intestinal Disorders Tissue Engineering and Cell Models Core for training, advice, and support in organoid culture and monolayer differentiation. We thank the University of Chicago Duchossois Family Institute Symbiotic Bacterial Strain Bank for providing the *Prevotella copri* strain and for assistance in preparing the microbe for the spike-in experiment. Figures in this manuscript were created and/or formatted using Biorender.com. This work was supported by NIH grants F30GM151855 to SN, R01GM109215 to FL and RPR, and R35GM128716 to RB.

## Supplemental Information

### Supplemental Figures

**Supplemental Figure 1.**
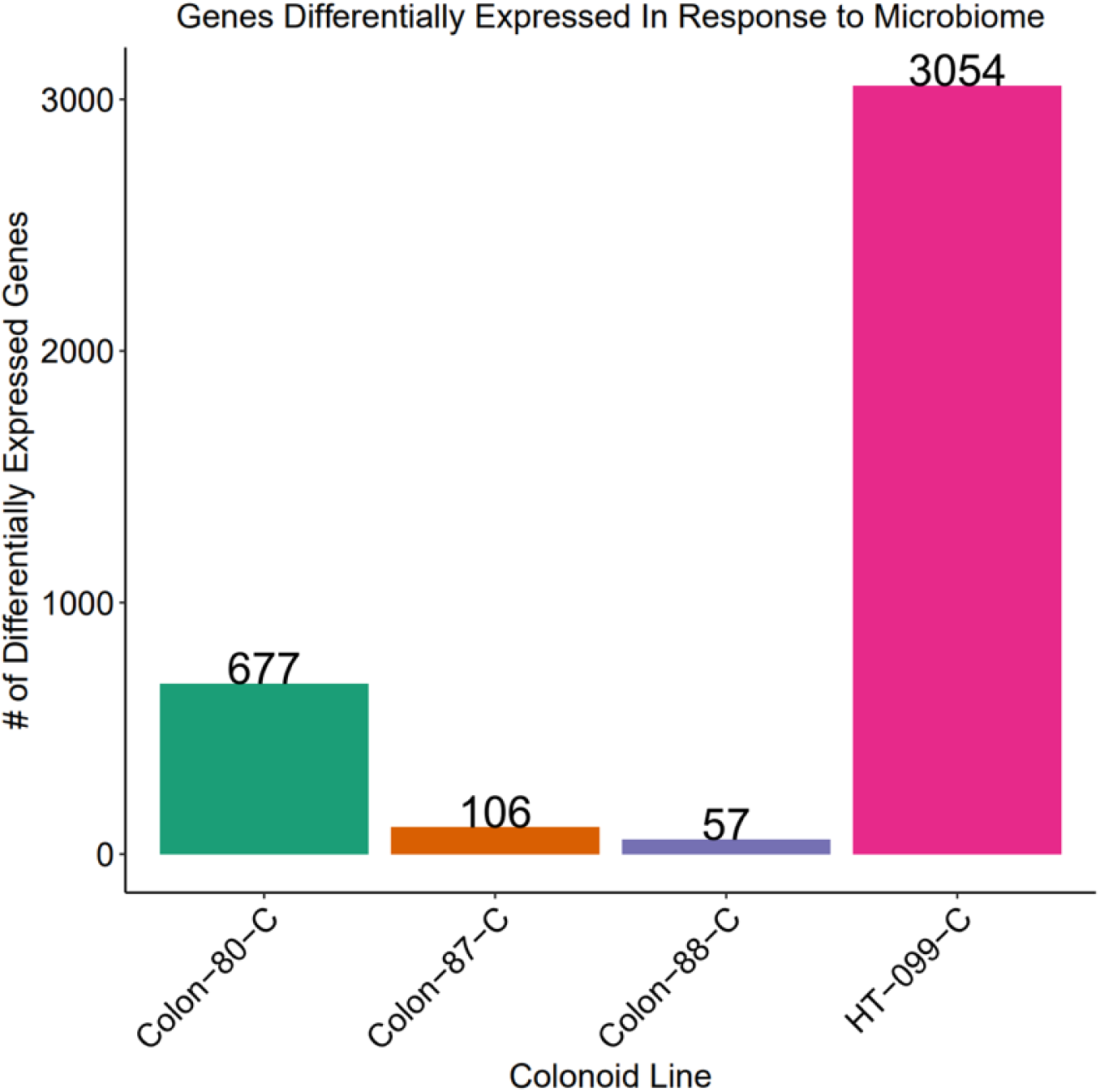
Host transcriptional response to microbiome exposure varies substantially between individuals. Barplot with number of significantly differentially expressed genes in response to microbiome exposure for the four responding colonoid lines (FDR < 10%).

**Supplemental Figure 2.**
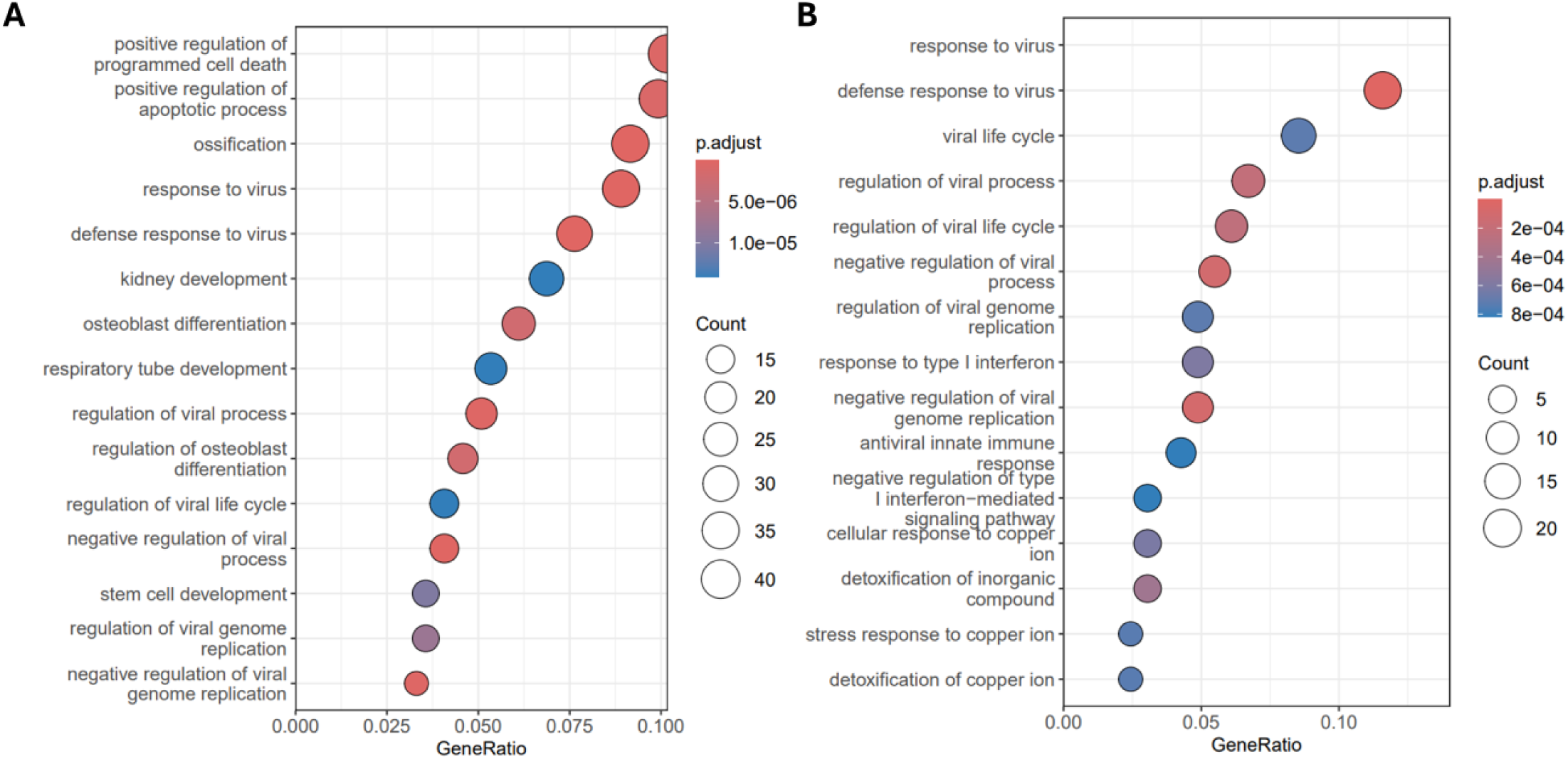
Urban and Rural microbiome responses are both enriched for downregulation of host viral response biological processes. Dotplots of top 15 enriched Gene Ontology biological processes terms for genes downregulated in response to A) Urban microbiome exposure or B) Rural microbiome exposure (FDR < 10%). The Y-axis is the GO term and X-axis is the GeneRatio, the ratio of genes associated with the GO term in significant differentially expressed genes over all differentially expressed genes. The size of the dot corresponds to the number of significant differentially expressed genes associated with the GO term and the color of the dot indicates the BH p-adjust value for the enrichment test.

**Supplemental Figure 3.**
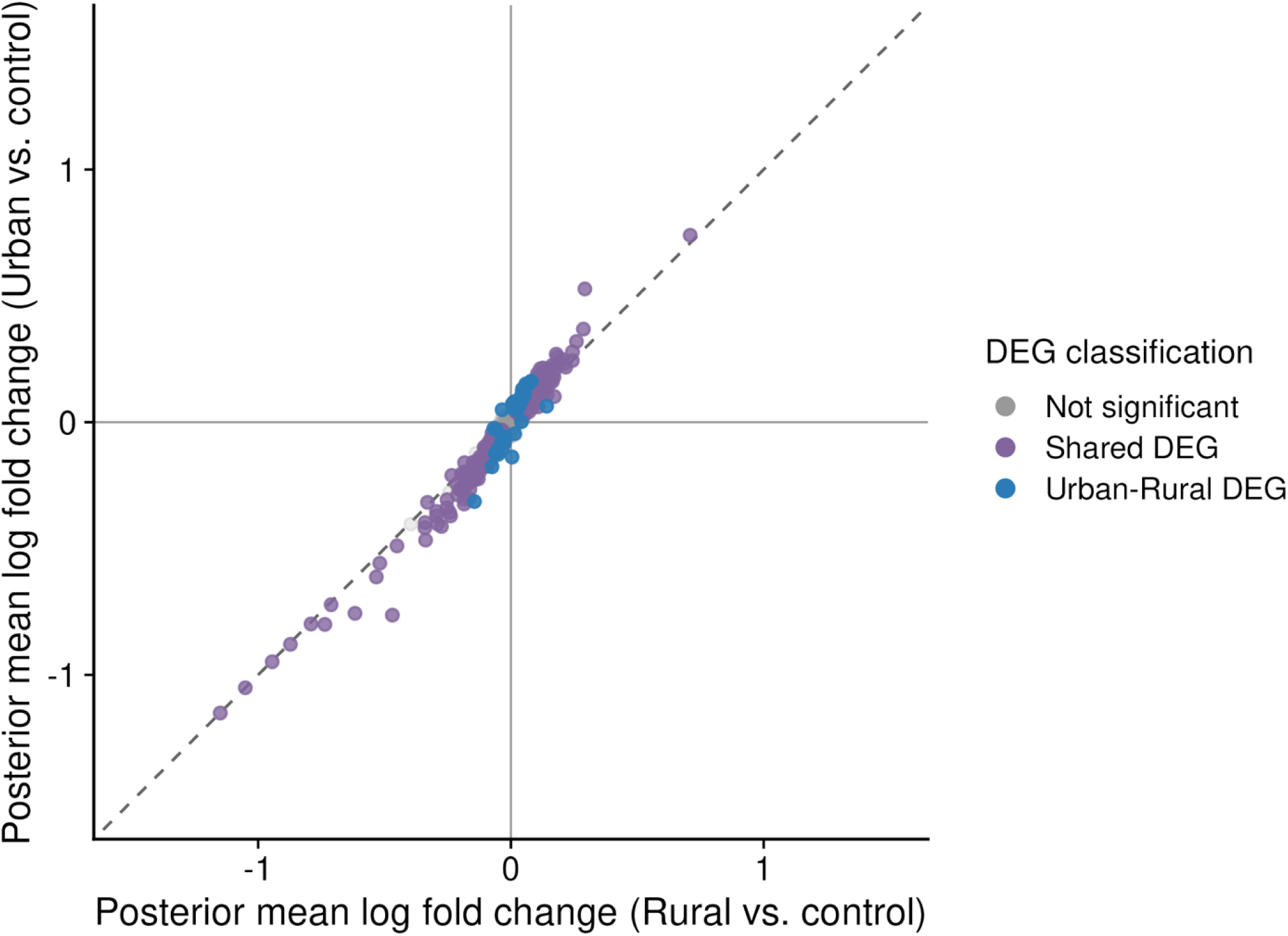
Scatterplot of posterior mean log fold changes in gene expression estimated by MASH for Urban microbiome versus control (y-axis) and Rural Microbiome versus control (x-axis) for all genes tested. Genes were classified as shared DEGs (purple), Urban-Rural DEGs (blue), or not significant (gray). The dashed gray line indicates the line of equality (y = x).

**Supplemental Figure 4.**
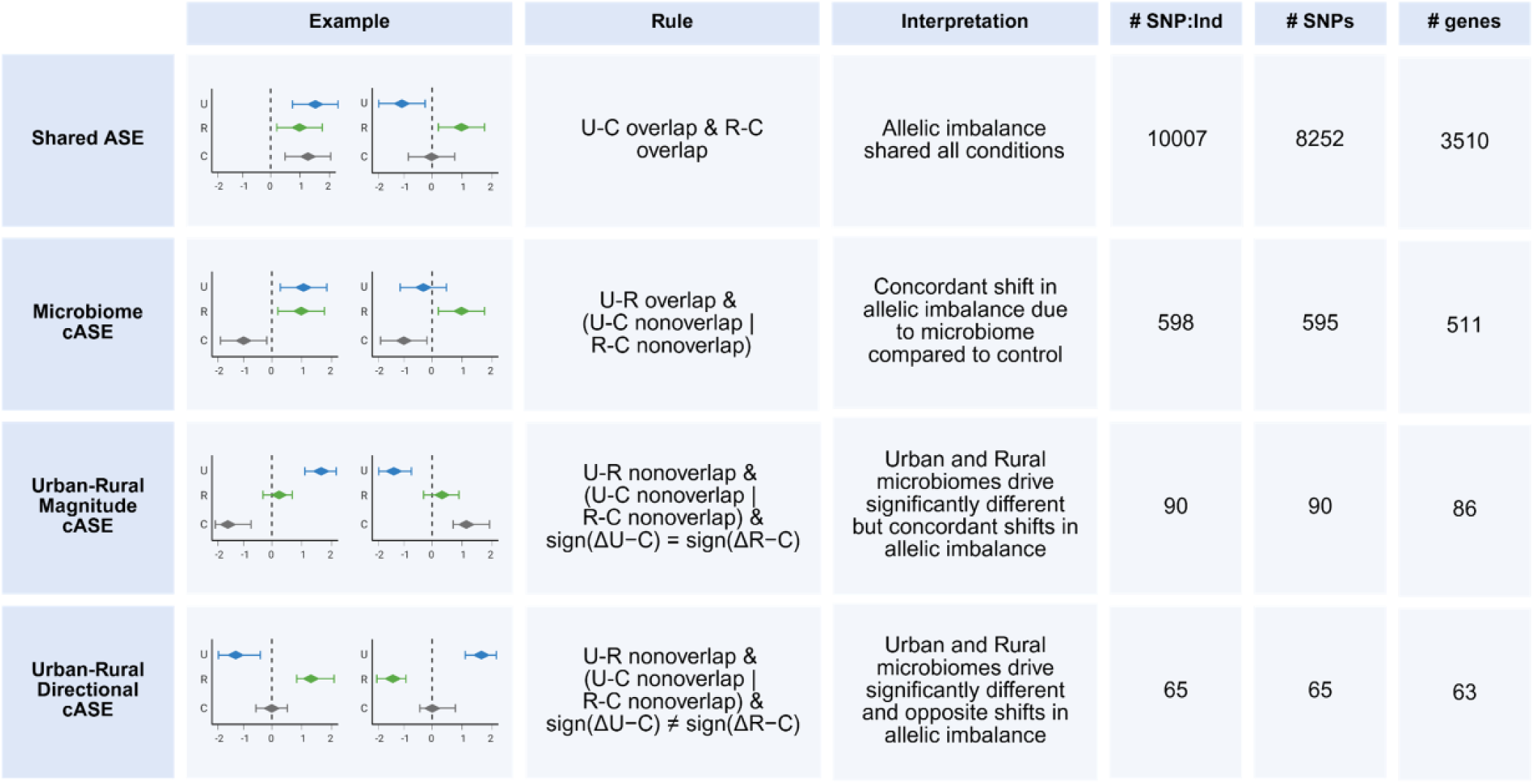
ASE classification. SNPs with ASE within a condition and individual are defined as having lfsr < 0.05 & |β| > 0.1. Union of SNP:Ind pairs across conditions were tested for shared and conditional ASE by testing for overlap of 95% credible intervals around posterior means generated by sampling the posterior distributions. Here we define four major classes of shared ASE/cASE defined by the rules as well as the interpretation of the overlap.

**Supplemental Figure 5.**
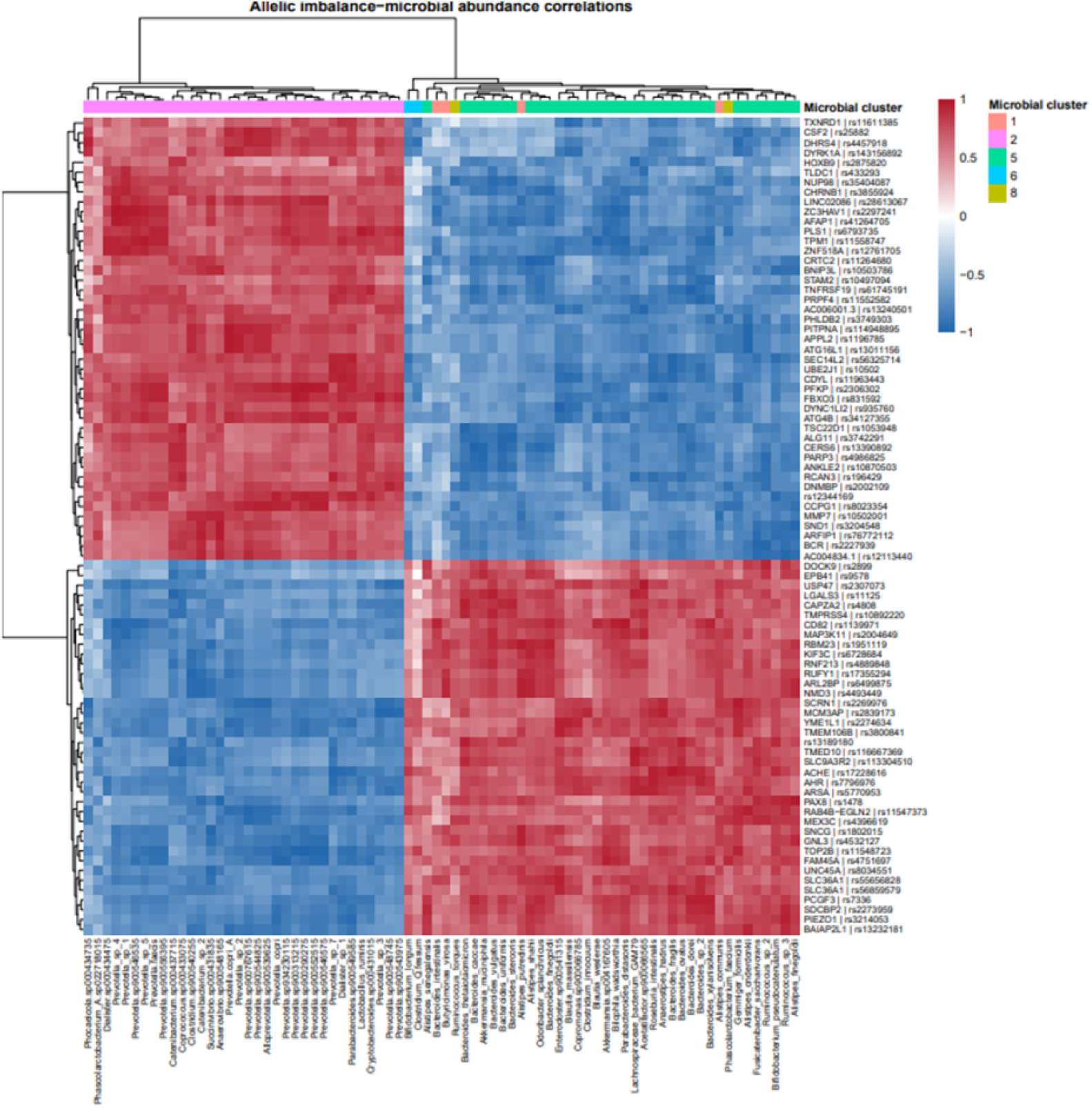
Correlation structure of Urban Rural cASE loci allelic imbalance-taxon associations. Heatmap of Spearman correlations between allelic imbalance and microbial taxon abundance for urban-rural specific cASE loci with at least one significant taxon association (FDR < 0.05). Rows represent SNP:donor pairs and columns represent microbial taxa, with colors indicating the Spearman correlation. Rows and columns were hierarchically clustered based on correlation profiles. The top annotation indicates microbial taxon clusters defined by abundance correlations across the eight microbiome samples.

**Supplemental Figure 6.**
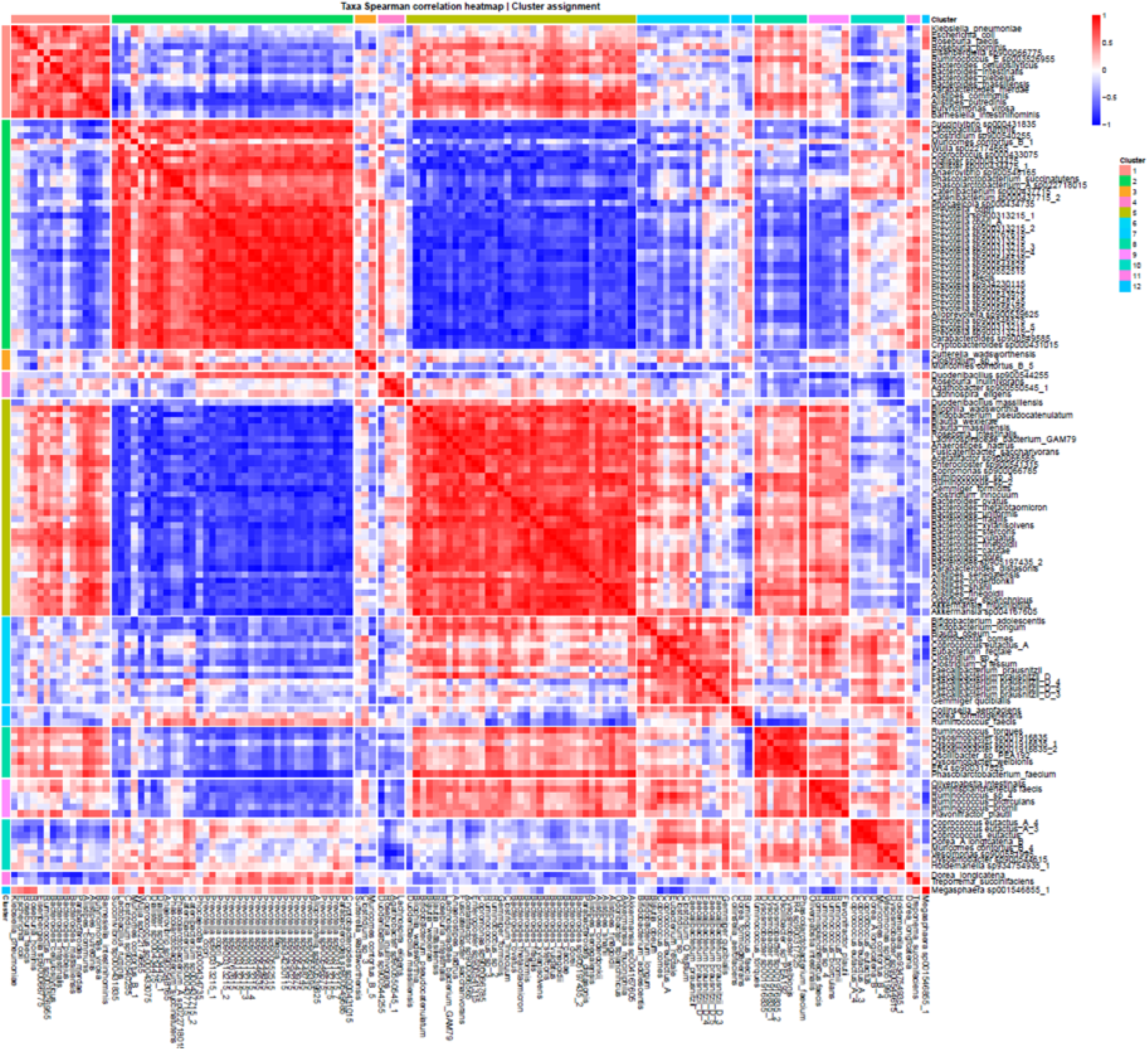
Taxon-taxon co-abundance correlation structure across the 8 microbiome samples. Heatmap of pairwise Spearman correlations (ρ) between CLR-transformed abundances of 136 filtered taxa across the 8 microbiome samples used in this study. Taxa are ordered by hierarchical clustering (average linkage; distance = 1 − ρ), and the colored bar (right) indicates assignment to one of 12 clusters.

**Supplemental Figure 7.**
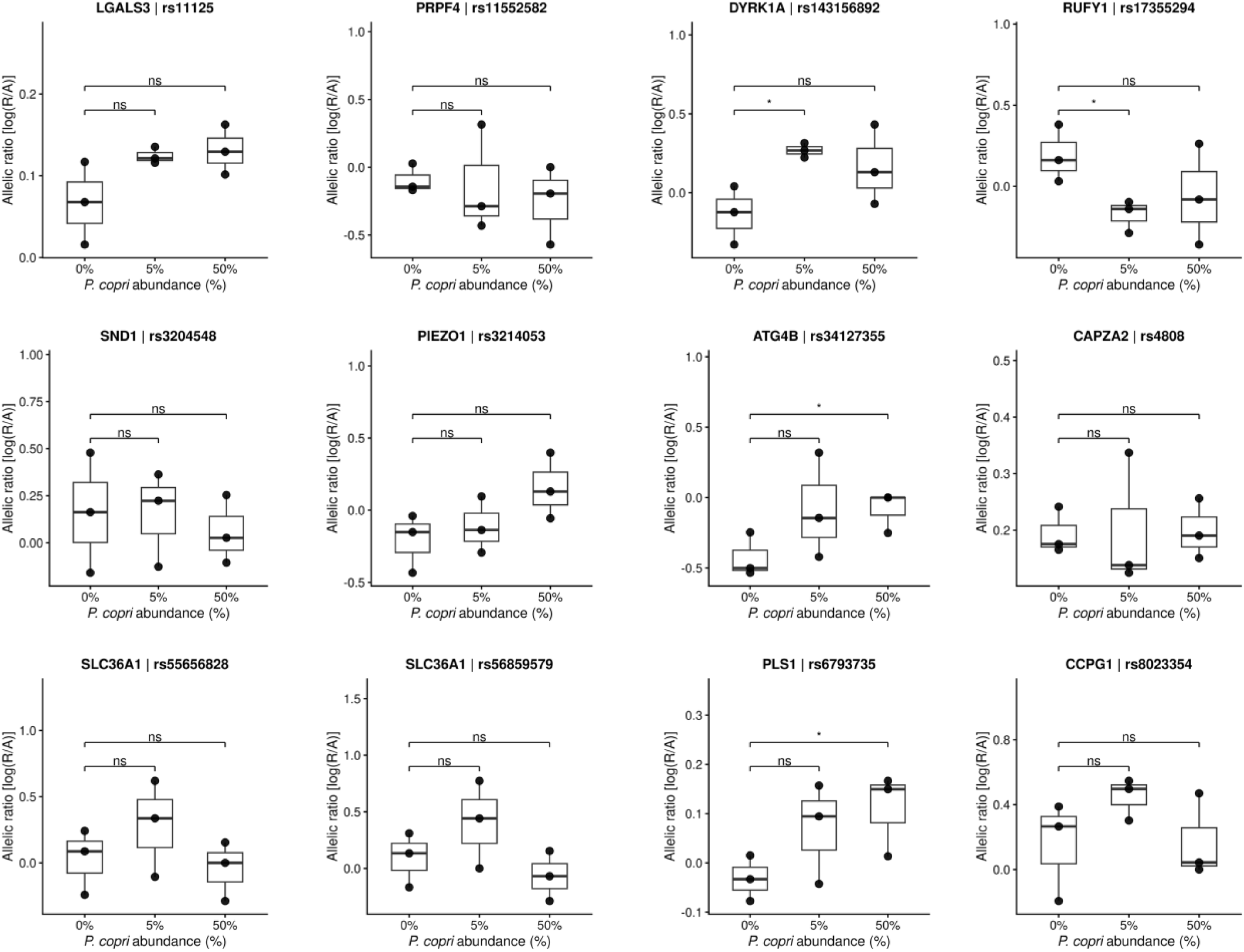
Pairwise comparisons between baseline (0%) and either 5% or 50% for loci with *Prevotella* associations that are present in the spike in experimental data. Y-axis is allelic ratio following *P. copri* MSK.21.56 spike-in at 0, 5 and 50% spike-in. Points represent individual experimental replicates (n = 3 per condition) and boxplots show distribution across replicates. Brackets indicate comparisons between 0% and either 5% or 50% *P. copri* spike in using one-sided Student’s t-tests, with the direction specified by the original *Prevotella*-allelic ratio association. P < 0.05; ns, not significant.

### Supplemental Tables

**Table 1 Microbial Sample Metadata. Contains metadata for the eight microbial samples used in this study.** Columns: Microbiome - Microbiome Sample ID; Urbanism - Rural or Urban lifestyle; Country - sample origin country; Lifestyle - donor lifestyle; Prevotella - Prevotella genus level relative abundance; Bacteroides - Bacteroides genus level relative abundance; Firmicutes - Firmicutes phylum level relative abundance

**Table 2 Host RNA Seq Sample Information and Sequencing Depth.** Contains sample metadata and sequencing information for each RNA-seq sample used in this study. Columns: Sample ID; Colonoid Line/Individual; Treatment (control or microbiome); Microbiome Sample Treatment ID; Total number of raw sequencing reads; Number of reads after quality control; Number of reads after quality control and removal of PCR duplicates; Fraction of raw reads remaining after quality control and deduplication; Fraction of clean reads mapping to exons

**Tables 3 - 6 DESeq2 Differential Gene Expression Results for Microbiome versus Control for individual colonoid lines.** Results of DESeq2 analysis for microbiome treated samples versus control for each colonoid line used. Columns: Gene - ENSEMBL ID; baseMean; log2foldchange; lfcSE - standard error of log 2 fold change; stat - test statistic; pvalue; padj - calculated using BH procedure; Colonoid_line; contrast - analysis

**Table 7 Meta-analysis of Microbiome versus Control Differential Gene Expression Results.** Colnames: Gene - ENSEMBL ID; baseMeanHat - meta-analyzed baseMean; beta - meta-analyzed log2foldchange; stderr - meta-analysis standard error; pval - meta-analysis pvalue; Qval - adjusted p-value using Storey’s q-value; contrast - analysis

**Tables 8 - 11 DESeq2 Differential Gene Expression Results for Urban Microbiomes versus Control for individual colonoid lines. Columns as in Tables 3 - 6.**

**Table 12 Meta-analysis of Urban Microbiome versus Control Differential Gene Expression Results. Columns as in Table 7**.

**Tables 13 - 16 DESeq2 Differential Gene Expression Results for Rural Microbiomes versus Control for individual colonoid lines. Columns as in Tables 3 - 6.**

**Table 17 Meta-analysis of Rural Microbiome versus Control Differential Gene Expression Results. Columns as in Table 7**.

**Tables 18 - 21 DESeq2 Differential Gene Expression Results for Urban versus Rural Microbiomes for individual colonoid lines. Columns as in Tables 3 - 6.**

**Table 22 Meta-analysis of Urban versus Rural Microbiome Differential Gene Expression Results. Columns as in Table 7**.

**Table 23 Multivariate Adaptive Shrinkage (MASH) Analysis of Urban and Rural Microbiome Transcriptional Responses.** Columns: Gene - ENSEMBL ID; Rural_lfsr - Rural Microbiome condition local false sign rate (lfsr); Urban_lfsr - Urban Microbiome condition lfsr; Rural_Posterior_Mean - posterior mean log2 fold change of Rural microbiome response; Urban_Posterior_Mean - posterior mean log2 fold change of Urban microbiome response; DEG_class - classification of gene based on lfsr significance and ratio of posterior mean log2 fold change effects

**Table 24 GO Biological Process Over-Representation Analysis (ORA) of Genes Upregulated in Response to Microbiome Treatment versus Control.** Colnames: ID - Gene Ontology (GO) term identifier; Description - GO term name; GeneRatio - number of upregulated genes annotated to the term / number of upregulated genes with GO annotations; BgRatio - number of background genes annotated to the term / number of background genes with GO annotations; FoldEnrichment - enrichment of the term among upregulated genes relative to background (GeneRatio / BgRatio); pvalue - hypergeometric test p-value; p.adjust - Benjamini-Hochberg adjusted p-value; Count - number of upregulated genes annotated to the term; geneID - gene symbols of the upregulated genes annotated to the term, separated by “/”.

**Table 25 GO Biological Process ORA of Genes Downregulated in Response to Microbiome Treatment versus Control.** Columns as in Table 24, with “downregulated” in place of “upregulated”.

**Table 26 GO Biological Process ORA of Genes Upregulated in Urban versus Control.** Columns as in Table 24.

**Table 27 GO Biological Process ORA of Genes Downregulated in Urban versus Control.** Columns as in Table 24, with “downregulated” in place of “upregulated”.

**Table 28 GO Biological Process ORA of Genes Upregulated in Rural versus Control.** Columns as in Table 24.

**Table 29 GO Biological Process ORA of Genes Downregulated in Rural versus Control.** Columns as in Table 24, with “downregulated” in place of “upregulated”.

**Table 30 Allelic Counts for Each SNP:Individual Pair Across All Replicates.** Colnames: Sample_ID - sample (replicate) identifier; identifier - unique SNP:individual identifier, formatted as rsID:individual; chr - chromosome; pos - genomic position of the SNP; rsID - dbSNP reference SNP identifier; ref - reference allele; alt - alternate allele; refCount - number of reads supporting the reference allele; altCount - number of reads supporting the alternate allele; total - total read count at the SNP (refCount + altCount)

**Table 31 Conditional Allele-Specific Expression (cASE) in Control, Urban, and Rural Conditions from Joint MASH Analysis.** Colnames: identifier - unique SNP:individual identifier, formatted as rsID:individual; rsID - dbSNP reference SNP identifier; individual - individual (donor) identifier; ensg - ENSEMBL ID of the gene containing the SNP; symbol - gene symbol; gene_class - gene biotype; pm_control; pm_urban; pm_rural - MASH posterior mean allelic ratio in the control, urban, and rural conditions; lfsr_control; lfsr_urban; lfsr_rural - MASH local false sign rate in each condition; lower_control; upper_control; lower_urban; upper_urban; lower_rural; upper_rural - lower and upper bounds of the 95% highest posterior density (HPD) interval of the allelic ratio in each condition; ase_control; ase_urban; ase_rural - whether the SNP showed significant allele-specific expression (allelic imbalance) in each condition (TRUE/FALSE); ase_pattern - conditions in which ASE was detected (e.g., control_urban_rural); delta_urban_control - pm_urban minus pm_control; delta_rural_control - pm_rural minus pm_control; delta_urban_rural - pm_urban minus pm_rural; overlap_class - classification of the SNP-donor pair by whether HPD intervals overlap between conditions (e.g., shared_all); bio_class_final - ASE class of the SNP-donor pair: shared_ASE - overlapping 95% HPD intervals between Urban and control and between Rural and control, indicating consistent allelic imbalance across all conditions; shared_microbiome_cASE - i.e. Microbiome cASE, non-overlapping HPD intervals between at least one microbiome condition and control, with overlapping Urban and Rural HPD intervals, indicating a shared microbiome-associated shift in allelic imbalance relative to control; urban_rural_cASE magnitude_diff - non-overlapping Urban and Rural HPD intervals and non-overlapping HPD intervals between at least one microbiome condition and control, with concordant Urban and Rural effects relative to control (sign(pmU - pmC) = sign(pmR - pmC), where pmU, pmR, and pmC are the posterior mean allelic ratios for Urban, Rural, and control); urban_rural_cASE opposite_direction - the same criteria as urban_rural_cASE magnitude_diff, but with discordant Urban and Rural effects relative to control

**Table 32 GO Biological Process ORA of Genes Containing SNPs in Each ASE Class.** Colnames: bio_class_final - ASE class of the genes tested (shared_ASE, shared_microbiome_cASE, urban_rural_cASE magnitude_diff, urban_rural_cASE opposite_direction, as defined in Table 31; all_urban_rural_cASE - magnitude_diff and opposite_direction combined; all_cASE - shared_microbiome_cASE and urban_rural_cASE combined); ID - GO term identifier; Description - GO term name; GeneRatio - number of genes in the ASE class annotated to the term / number of genes in the ASE class with GO annotations; BgRatio - number of background genes annotated to the term / number of background genes with GO annotations; pvalue - hypergeometric test p-value; p.adjust - Benjamini-Hochberg adjusted p-value; Count - number of genes in the ASE class annotated to the term; geneID - gene symbols of the genes in the ASE class annotated to the term, separated by “/”; n_fg_ensg - number of ENSEMBL gene IDs in the ASE class (foreground) gene set. The background was all genes with testable SNPs.

**Table 33 KEGG Pathway ORA of Genes Containing SNPs in Each ASE Class.** Columns as in Table 32 (ASE classes as defined in Table 31 and Table 32), with the following differences: ID - KEGG pathway identifier; Description - KEGG pathway name; GeneRatio and BgRatio - based on KEGG annotations; geneID - gene symbols of the genes annotated to the pathway; category - KEGG pathway category; subcategory - KEGG pathway subcategory; n_fg_entrez - number of genes in the foreground set with Entrez IDs. The background was all genes with testable SNPs.

**Table 34 Spearman Correlations Between Allelic Ratio at Urban-Rural cASE Loci and CLR-Transformed Abundance of Microbial Taxa.** Colnames: identifier - unique SNP:individual identifier, formatted as rsID:individual; taxon - microbial taxon; n - number of microbiome treatments with allelic ratio measurements; rho - Spearman correlation coefficient between allelic ratio (log(Ref/Alt)) and CLR abundance of the taxon across microbiome treatments; pval - Spearman correlation p-value; padj - Benjamini-Hochberg adjusted p-value across all 11,832 tests

**Table 35 Prevotella Spike-In Validation Sample Information, Sequencing Quality Control, and Total Reads.** Colnames: Sample_Name - sample identifier; Colonoid_Line - colonoid line (donor) identifier; microbe_ID - microbiome treatment identifier (FMB028 - microbiome without Prevotella spike-in; prevotella_5 - Prevotella spike-in at 5%; prevotella_50 - Prevotella spike-in at 50%); Replicate_ID - replicate identifier; prevotella_dose - Prevotella spike-in dose (0, 5, or 50); TotalReads - total number of sequencing reads; Quality - number of reads passing quality control; Clean - number of reads after quality control and removal of PCR duplicates; Fraction_Clean_Reads - fraction of reads remaining after quality control and removal of PCR duplicates; Fraction_Exon_reads - fraction of clean reads mapping to exons

**Table 36 Prevotella Spike-In Validation at Urban-Rural cASE Loci.** Colnames: identifier - unique SNP:individual identifier, formatted as rsID:individual; n_prevotella_taxa - number of Prevotella taxa whose abundance significantly correlated with allelic ratio at the locus in the primary analysis; prevotella_taxa - names of those taxa, separated by “;”; n_original_positive; n_original_negative - number of those taxa with a positive or negative correlation in the primary analysis; original_directions_consistent - whether all significant correlations at the locus had the same direction (TRUE/FALSE); dose_slope - slope of the linear regression of allelic ratio (log(Ref/Alt)) on Prevotella spike-in proportion (0, 0.05, 0.50); effect_0v5; effect_0v50 - mean allelic ratio at 5% or 50% spike-in minus mean at 0%; effect_presence - mean allelic ratio in spike-in samples (5% and 50%) minus mean in samples without spike-in (0%); predicted_direction - direction of allelic ratio change expected with Prevotella spike-in, based on the sign of the primary correlations (increase or decrease); dose_p; p_0v5; p_0v50; p_presence - one-sided p-values (dose-response slope; Student’s t-test for 0% versus 5%, 0% versus 50%, and spike-in versus no spike-in) in the predicted direction; concordant_dose; concordant_0v5; concordant_0v50; concordant_presence - whether the observed direction matches predicted_direction (TRUE/FALSE); sig_dose; sig_0v5; sig_0v50; sig_presence - whether the corresponding p-value is < 0.05 (TRUE/FALSE)

**Table 37 Conditional Allele-Specific Expression (cASE) Genes with Significant S-PrediXcan Associations to GWAS Traits.** Colnames: gene_name - gene symbol; ensg - ENSEMBL gene ID; cASE_class - cASE class of the gene (shared_microbiome_cASE, urban_rural_cASE magnitude_diff, or urban_rural_cASE opposite_direction); trait - GWAS trait; tissue - GTEx tissue of the prediction model (Colon_Sigmoid or Colon_Transverse); zscore - S-PrediXcan z-score of the gene-trait association; pvalue - S-PrediXcan p-value; best_gwas_p - smallest GWAS p-value among the variants in the gene’s prediction model. Only gene-trait-tissue associations with S-PrediXcan p < 0.05 are shown.

